# CRISPR-Hybrid: A CRISPR-Mediated Intracellular Directed Evolution Platform for RNA Aptamers

**DOI:** 10.1101/2023.08.29.555185

**Authors:** Qiwen Su-Tobon, Jiayi Fan, Kevin Feeney, Hongyuan Ren, Patrick Autissier, Peiyi Wang, Yingzi Huang, Jia Niu

## Abstract

Recent advances in gene editing and precise regulation of gene expression based on CRISPR technologies have provided powerful tools for the understanding and manipulation of gene functions. Fusing RNA aptamers to the sgRNA of CRISPR can recruit cognate RNA-binding protein (RBP) effectors to target genomic sites, and the expression of sgRNA containing different RNA aptamers permit simultaneous multiplexed and multifunctional gene regulations. Here, we report an intracellular directed evolution platform for RNA aptamers against intracellularly expressed RBPs. We optimized a bacterial CRISPR-hybrid system coupled with FACS, and identified novel high affinity RNA aptamers orthogonal to existing aptamer-RBP pairs. Application of orthogonal aptamer-RBP pairs in multiplexed CRISPR allowed effective simultaneous transcriptional activation and repression of endogenous genes in mammalian cells.

## Introduction

Genome editors use programmable sequence-specific nucleases^1–3^ to generate DNA double-strand breaks (DSBs) at predetermined sites and introduce DNA insertions/deletions by non-homologous end joining, homology-directed repair, or microhomology-mediated end joining^4–7^. The emergence of CRISPR (Clustered Regularly Interspaced Short Palindromic Repeats)-Cas systems has accelerated the development of DSB-dependent and DSB-independent genome editing technologies^8–14^. CRISPR consists of two components: a CRISPR-associated nuclease (Cas, *e.g.*, Cas9) and an artificial single guide RNA (sgRNA). Repurposing of CRISPR by inactivating the nuclease domain (*e.g.*, catalytically dead Cas9, dCas9)^15,16^ and covalently tethering functional effectors to dCas9 have enabled precise gene modifications, such as base editors^17–21and^ prime editors^9,22^, epigenetic editing^23–27^ such as methylation^28–31^ and acetylation^32^, and gene expression regulation^33–36^ including activation^37–40^ and repression^41,42^. Besides modifying dCas9, the sgRNA scaffold can also be adapted to accommodate incorporations of RNA aptamers^43^ to directly recruit RNA-binding proteins (RBP) to the target site^38,44–47^. Multiplexed gene regulation events in the same cell can be coordinated by expressing sgRNA scaffolds carrying orthogonal aptamers and recruiting different numbers and types of effectors to the target region^44,48^. Despite the great potential, there have been only a limited set of mutually orthogonal aptamer-RBP pairs that function intracellularly, posing a significant constraint on the utility of multiplexed CRISPR for simultaneous multifunctional genome editing and transcriptional modulation.

RNA aptamers with specificity and high affinity are traditionally generated by a repetitive *in vitro* selection process termed SELEX (systematic evolution of ligands by exponential amplification)^49,50^. Classic SELEX protocols typically consist of four steps: target incubation, wash-off unbound sequences, elution, and amplification of the bound sequences. Although SELEX is well-established, it is considered as a time-consuming and labor-intensive method. Most importantly, *in vitro* selected aptamers often fail to bind to targets in their natural biological environment. To overcome this limitation, *in vivo* selection strategies have been proposed as an alternative to SELEX for discover aptamers for intracellular applications. For example, Liu et al. reported a yeast three-hybrid system (Y3H)-based selection strategy to identify RNA and protein binding partners intracellularly^51–54^. We took inspirations from Y3H and aptamer-mediated CRISPR-dCas9 systems to develop CRISPR-hybrid to intracellularly evolve functional RNA aptamers as part of the sgRNA. In a CRISPR-hybrid system, the dCas9 binds to a promoter region upstream to a gene encoding a fluorescent reporter. The sgRNA scaffold is extended with a randomized RNA aptamer pool sequence, which is challenged to recruit the target protein that is fused to a transcriptional activator for reporter expression. Our approach implements fluorescent-activated cell-sorting (FACS) to simultaneously isolate cell population carrying the functional aptamer from unbound species. Using this method, we identified a new RNA aptamer-RBP pair orthogonal to all existing pairs when expressed in bacteria and mammalian cells. We demonstrated that simultaneous transcription activation and repression of different genes could be achieved multiplexed using multiplex sgRNAs carrying orthogonal aptamer-RBP pairs. It is noteworthy that parallel to our work, Zhang *et al.* recently reported another CRISPR/Cas-based screening system for RNA aptamers targeting the extracellular receptor binding domain of the spike glycoprotein of SARS-CoV-2^55^, but this study fell short of identifying aptamers for intracellular targets.

## Results

### Design and validation of the CRISPR-hybrid system

The CRISPR-hybrid system consists of four components: a dCas9 protein, a sgRNA scaffold extended with one copy of aptamer to be evolved (hereafter as the sgRNA-aptamer library chimera), an RBP-transcriptional activator fusion protein, and a gene encoding a selection marker or a fluorescent reporter (**Fig. 1a**). Our platform relies on the ability of sgRNA-aptamer library chimera to recruit target RBP to be proximal to the gene encoding the selection marker/fluorescent reporter such that its transcription is activated. We distributed these components into three plasmids (SP, AP, and RP) carrying orthogonal origins of replication and antibiotic resistance cassettes such that the individual components of the selection system can be independently varied.

**Figure 1.**
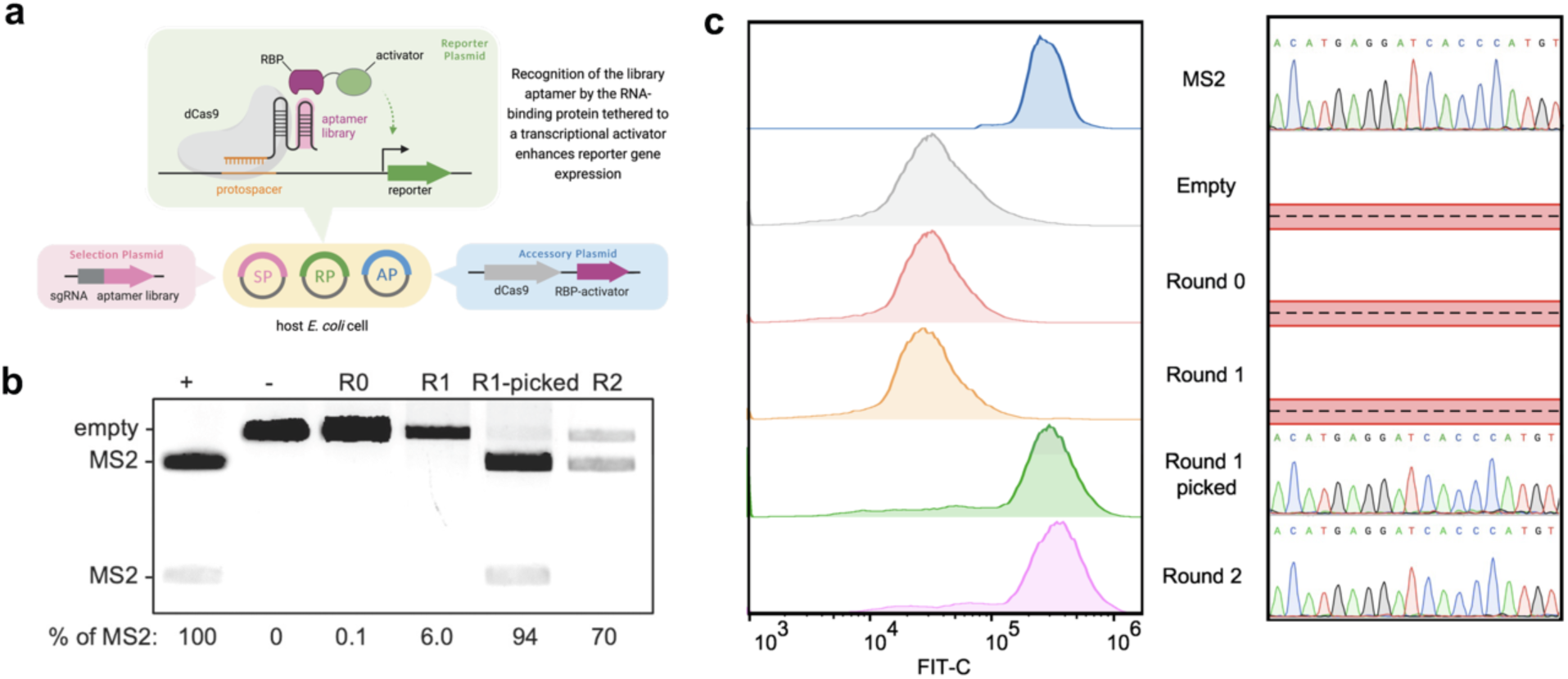
Intracellular aptamer selection using CRISPR-hybrid. **a**, Overview of the selection strategy. **b,** Gene compositions before and after mock selection were compared with *ScaI* and *AleI* digestion, showing >940-fold enrichment of MS2. **c**, FACS histogram overlays of mock populations (left), showing significant enrichment of cell population with the *MS2* gene after two rounds, and Sanger Sequencing of selection plasmids (right) in each round.

To maximize the sensitivity of the transcriptional activation, we used a known MS2-bacteriophage coat protein (MCP) aptamer-RBP pair and the *sfGFP* gene that encodes the super-fold GFP reporter to optimize parameters including (1) sgRNA binding site, (2) sgRNA binding site-promoter distance, (3) target RBP-transcription activator linker length, (4) protein expression level, (5) reporter gene ribosome-binding site (**Suppl. Fig 1a-e**). After initial optimizations, a 5.4-fold transcription activation was achieved for the optimized construct that contains the *Escherichia coli* RNA polymerase omega subunit (RpoZ) as the transcriptional activator, a G6’ sgRNA binding at 16 base pairs upstream of -35 signal of the J23107 promoter, and promoter from *Streptococcus pyogenes* to drive dCas9 transcription.

Next, we sought to couple transcriptional activation with a reporter suitable for intracellular evolution of aptamers, and antibiotic-based survival selection was first attempted. Using the MS2-MCP pair, we screened various antibiotic resistance reporters and identified a kanamycin reporter in which cells that contain the CRISPR-hybrid construct except the aptamer survived up to 1.0 mg/ml kanamycin while introducing the MS2 aptamer to these cells led to survival up to 1.5 mg/ml kanamycin (**Suppl. Fig 2a**). To test the efficacy of this system, we designed and executed a mock selection to enrich a MS2-incorporated sgRNA (“MS2-sgRNA”) from a sgRNA scaffold lacking any aptamers (“empty sgRNA”) at 1:1000 ratio against kanamycin. In one round, we observed ≥403-fold enrichment of the *MS2* gene (**Suppl. Fig. 2b**). However, subsequent rounds of selection with identical or higher concentrations of kanamycin did not result in further enrichment of the *MS2* gene. We attributed the lack of productive enrichment to the low degree of transcription activation and the high background caused by RpoZ-independent transcription. These results suggest that the dynamic range of the RpoZ-based construct is inadequate for an effective survival-based aptamer selection system.

To circumvent this issue, we changed the RBP-fused transcriptional activator from RpoZ to SoxS_R93A_, which has been shown to enhance transcription activation in CRISPR constructs^45^ (**Suppl. Fig. 3a**). Indeed, this modification resulted in a 20-fold increase in sfGFP expression after the sgRNA binding site and RNA polymerase-promoter affinity were further optimized (**Suppl. Fig. 3b, c**). We also confirmed that the enhanced reporter activation arose from the specific recruitment of MCP-SoxS by the MS2 aptamer (**Suppl. Fig. 3d**). Consistent with the improved reporter activation, cells consisting of the SoxS-based construct showed resistance to up to 5 mg/mL of kanamycin (**Suppl. Fig. 4a**). We repeated mock selection by challenging a 1000:1 mixture of cells containing empty sgRNA and MS2-sgRNA to activate an upstream anti-kanamycin selection marker and a downstream sfGFP reporter. Despite higher enrichment (≥583-fold) of the *MS2* gene was observed (**Suppl. Fig. 4b**), the library still predominantly consisted of inactive variants. These results indicated that the antibiotics survival selection approach may become limited for intracellular evolution of aptamers due to the high background activation.

During the antibiotic selection, we noticed that the survived cells can be distinguished as two separate populations on sfGFP fluorescence histograms (**Suppl. Fig. 4c**). Cells containing MS2 exhibited higher levels of fluorescence, while cells lacking the aptamer exhibited only background levels of fluorescence. Inspired by this finding, we explored an alternative method of fluorescence-based selection using fluorescence-activated cell sorting (FACS) by coupling transcription activation with the expression of a sfGFP gene cassette. We carried out a mock selection to enrich the *MS2* gene from a 1:1000 mixture with the sgRNA-only cells to validate the FACS-based selection method. Cells were grown at 37°C for 17 hours for maximal sfGFP reporter gene expression (**Suppl. Fig. 5a**). The top 5 out of every 100,000 cells expressing the most fluorescence, were collected in at least 1 mL of rich media (**Suppl. Fig. 5b**). A total of 10^8^ cells were sorted in each round to isolate the most fluorescent cells. To avoid the potential survival bias caused by the phototoxicity to cells during FACS, the post-sorting cells were not directly amplified; instead, they were lysed after sorting and the DNA fragments encoding the enriched sgRNA-aptamer chimera were PCR amplified, followed by assembly with selection plasmid backbone and reintroduced into selection strain carrying appropriate accessory and reporter plasmids for subsequent rounds (**Suppl. Fig. 5c-d**). Selection plasmids before and after the mock selection were digested with *Sac*I and *Ale*I, which specifically cleaves plasmid backbone and the *MS2* gene, respectively, suggesting that the *MS2* gene was successfully enriched by ≥940-fold after only two rounds of FACS-based mock selection (**Fig. 1b**). FACS histogram overlays showed the progress selection with an evident appearance of a second distinct cell population after two rounds (**Fig. 1c left**). Twenty colonies were sequenced in each round and the *MS2* gene was observed in the second population (**Fig. 1c right**). These results validated that our designed intracellular selection platform using FACS can strongly enrich active binders from mixtures that predominantly contain inactive or less active variants.

### Initial intracellular selection

We then sought to discover aptamers for intracellularly expressed proteins using the intracellular selection strategy. To this end, we generated an aptamer library derived from the MS2 aptamer by randomizing 11 nucleotides in the upper stem and loop regions, while keeping the lower stem of MS2 intact (**Supple Fig. 6a**). We reason that this design maintained the stability of the hairpin and could cause minimal disruption to sgRNA scaffold structures^46^. The theoretical diversity of the library is 4^11^ = 4.2 x 10^6^ unique sequences, and we obtained ∼5 x 10^8^ transformants, providing a diversity coverage greater than 100-fold. As a proof-of-principle. we first applied this library to the selection with the well-characterized target MCP. We screened ∼10^9^ cells on a single round of FACS. Roughly 25% of population exhibited higher fluorescence post sorting, and a second population emerged after only one round by setting the sorting gate to include top 7 cells out of every 100,000 cells (**Fig. 2a left**). It is noteworthy that a less stringent sorting gate (top 45 cells out of every 100,000 cells) failed to further enrich the high fluorescent population, suggesting that a tighter sorting gate is critical to provide sufficient selection pressure necessary for a successful round. After four rounds of selection, the Sanger Sequencing revealed the most abundant sequence to be the *MS2* gene, the consensus aptamer for MCP (**Fig. 2a right**).

**Figure 2.**
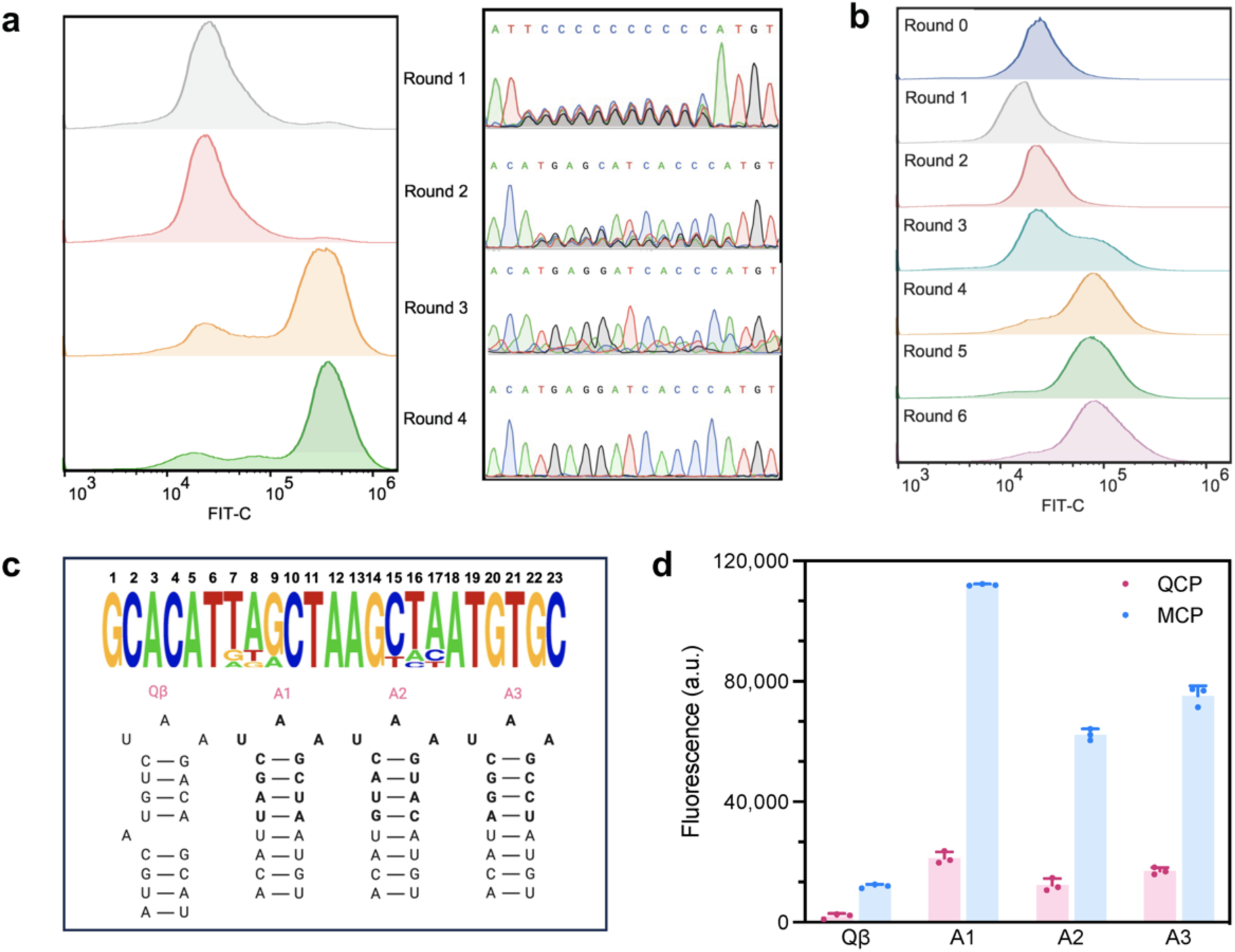
Initial intracellular selection of N11 aptamer library. **a**, FACS histogram overlays of four rounds selection for MCP. **b**, FACS histogram overlays of six rounds selection for QCP. **c,** Analysis of the 1% most-abundant sequences from round 6 (*n* = 15) is shown above, with the relative abundance of each base at each randomized position. The secondary structures of Qβ and the top three hits for QCP are shown below. **d,** GFP fluorescence measurements of aptamers binding to QCP and MCP. Values are mean ± s.d. (*n* = 3 independent replicates).

To evolve new RNA-RBP pairs, we next applied our selection system and library N11 to discover RNA aptamers for bacteriophage Qβ coat protein (QCP) (**Suppl. Fig. 6b**). QCP represents a unique RBP with a RNA binding interface distinct from that of MCP^56^. We first screened ∼10^9^ cells on FACS and collected top 7 cells out of every 100,000 cells each round. Enrichment was significantly evident after four rounds, with a distinct population displaying increased fluorescence (**Fig. 2b**). Further enrichments were modest in the next two rounds. We then subjected the recovered aptamer pools from each round to high-throughput DNA sequencing (HTS) analysis. Sequences are QC-filtered, counted for abundance, normalized to total number of reads per million (RPM) and ranked according to decreasing abundance. We compared the genotypic frequencies in each round of selection by plotting RPM values of one round against the immediate next round (**Suppl. Fig. 6c**). We observed a group of sequences that is shifted below the diagonal and a large group that is shifted notably upward, indicating successful aptamer evolution. These results matched with the fluorescence histograms displaying round-to-round enrichment (**Fig. 2b**). The scatter plots of round 5 and 6 showed a vast majority of the sequences clustered closely to the diagonal, suggesting these two populations are highly similar, as expected from observing their corresponding fluorescence histograms (**Suppl. Fig. 6b, Fig. 2c**). Analysis of the top 1% most-abundant sequences displayed an overrepresented sequence motif with a UAA loop and CG base pair in the first position of the upper stem, which resembled the previously reported aptamer Qβ for QCP^57^ (**Fig. 2c**). We analyzed three most-abundant hits (A1 to A3), and all were unique sequences containing UAA loop, ^1^CG base pair, and three different base pairs in the remaining upper stem region. Validation of A1 to A3 using a GFP reporter, including Qβ as reference, showed that selected aptamers induced stronger transcriptional activation than Qβ (**Fig. 2d**). These results confirmed that efficient binding to QCP requires a three-nucleotide loop and four base pairs upper stem, but the sequence of the stem is not critical for QCP binding^57^ as A1 to A4 have altered upper stems.

Unexpectedly, A1 to A3 also exhibited strong binding preference for MCP (**Fig. 2d**). We hypothesize that affinity for MCP was most likely because these motifs contain the MS2 lower stem that was not randomized in the library. To dissect key sequence-activity relationships among evolved aptamers, we selected the most abundant sequence A1, and introduced nucleotide mutations in the -5 and -7 position of loop region to profile its specificity for QCP using a GFP reporter (**Suppl. Fig. 7a-b**). Reduced binding to both QCP and MCP were observed with all variants. The interactions between variant aptamers and the bacteriophage coat proteins were weaken likely due to loss of protein contacts in the binding pocket, which is consistent with previous findings that -5A is a crucial residue for QCP binding^56–58^. To examine the contribution of MS2 lower stem sequence to the specificity of A1, we systematically mutated each base pair to the remaining three possibilities. We hypothesize that slight modifications in the lower stem could possibly tune A1 selectivity while preserving binding affinity to QCP. Indeed, all the variants showed significant lower binding to MCP (**Suppl. Fig. 7c**). The binding affinity to QCP remained constant except for variants 1UA and 1GC, which shown drastically decrease in binding. Further rational mutations only led to modest improvements in specificity. Together, these results suggest that lower stem is critical for improving the specificity of our selected aptamer, and additional rounds of evolution, rather than rational mutation, is needed to further improve the specificity.

### Second intracellular selection

We used first-generation aptamer A1 as the starting point for a second intracellular selection toward QCP binding. Eight bases in the lower stem of A1 were randomized. Transformation of cells with this library produced 5×10^8^ transformants, ensuring a ∼10^4^-fold coverage of the library with a theoretical diversity of 6.5 x 10^4^ (**Suppl. Fig. 8a**). Four rounds of selections were completed using FACS (**Fig. 3a**). Sequence pools recovered from each round of sorting were analyzed with HTS. The sequences in Round 3 and 4 were distributed along the diagonal of the scattered plot (**Suppl. Fig. 8b**), suggesting that these two populations are highly similar. In contrast to the first library, an overrepresented sequence was not observed in the second selection (**Fig. 3b**). A large number of sequences were present in comparable abundances (**Suppl. Fig. 8b**); therefore, we analyzed sequences in the top 10 most-abundant hits of rounds 3 and 4, A5-A14, and observed enrichment of bulge bases in the first and second positions of the lower stem. Of the ten unique sequences, three contained single bulged nucleotide, and three contained a mismatched base pair on the first position of the lower stem. Three sequences contained a mismatched base pair on the second position (**Fig. 3b**). No sequence contained more than one mismatched base pair. We profiled the activity of A5 to A14 with QCP and MCP to see if they favor their respective target QCP over MCP. The specificity profile of all the motifs exhibited lower affinity to MCP while their binding to QCP is similar to A1 or slightly reduced (**Fig. 3c**). Notably, aptamer A9 exhibited significantly higher specificity for QCP versus MCP. Binding affinity characterized by surface plasmon resonance (SPR) confirmed that A9 binds strongly to QCP with a dissociation constant (*K*_d_) of 10.2 nM, compared to the weak affinity to MCP *K*_d_ of 95.1 nM (**Fig. 3d, Suppl. Fig. 9**). Extending A9 with the sgRNA scaffold further improved the affinity and specificity, with a *K*_d_ of 6.21 nM to QCP and 132 nM to MCP, respectively (**Fig. 3d, Suppl. Fig. 9**). These *in vitro* affinity results are consistent with the FACS-based fluorescent measurements of the cells expressing the sgRNA-A9 chimera, confirming that A9 is highly specific to QCP both *in vitro* and in cells.

**Figure 3.**
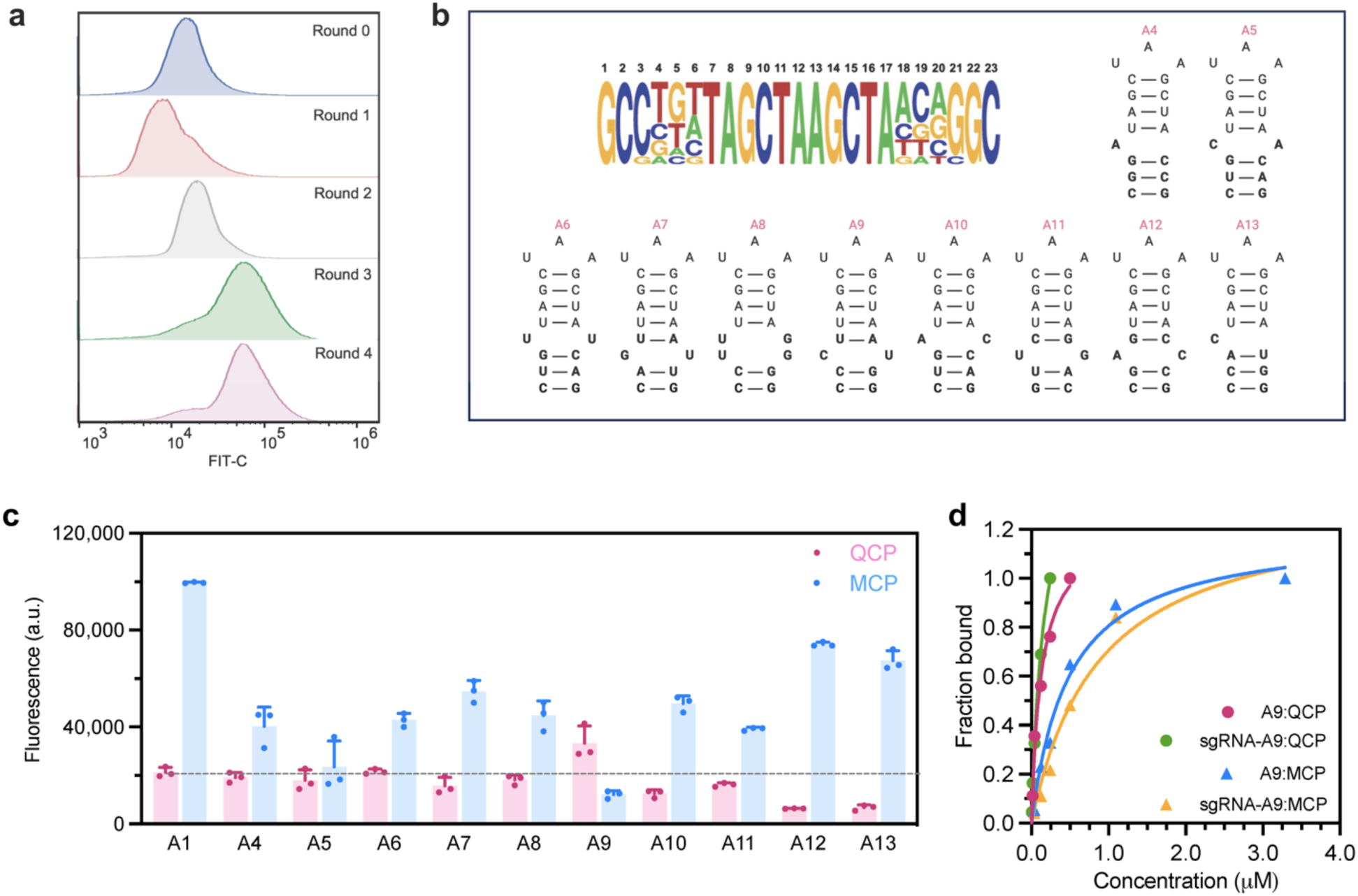
Second intracellular selection of N8 aptamer library. **a**, Overlaid histograms shows aptamer library was fully enriched after four rounds of FACS for QCP. **b**, Analysis of 1% most-abundant sequences from round 4 (n = 15) is shown, with the relative abundance of each base at each randomized position. The secondary structures of top ten hits for QCP are also shown. **c,** Fluorescence measurements of top ten aptamers binding to QCP and MCP. Values are mean ± s.d. (*n* = 3 independent replicates). **d**, SPR characterizations of A9 and sgRNA-A9 to QCP and MCP.

### Orthogonal RNA-protein recruitment in mammalian cells

Orthogonal aptamer-protein interactions in multiplexed CRISPR genome editing are critical for reducing crossbinding between components and minimizing off-target effects. To determine whether there is crossbinding between aptamers and non-cognate proteins in multiplexed CRISPR genome editing, we expressed three previously reported consensus aptamers (MS2, Qβ, and PP7) and A9 in HEK293T mammalian cells containing either MCP, QCP or bacteriophage PP7 coat protein^59^ (PCP) fusion proteins for luciferase reporter ([Fluc]) activation (**Fig. 4a**). No crossbinding was detected between PP7 and non-cognate proteins. While no crossbinding was found between MS2 and PCP, modest crossbinding was observed between MS2 and QCP. In contrast, significant crossbinding was found between Qβ and MCP and PCP, suggesting that this previously reported QCP aptamer is not orthogonal to other aptamer-RBP pairs when applied *in vivo*. Intriguingly, no crossbinding was observed between A9 aptamer and non-cognate proteins, indicating that this RNA aptamer enriched through our intracellular selection method exhibits high specificity for the cognate target in mammalian cells. In addition, A9 mediated robust [Fluc] activation with QCP over MCP by 135-fold (**Fig. 4b**). The strong activation of reporter gene by orthogonal modules (MS2-MCP, A9-QCP, and PP7-PCP) demonstrates the potential for simultaneous, locus-specific, independent transcriptional regulation of multiple genes.

**Figure 4.**
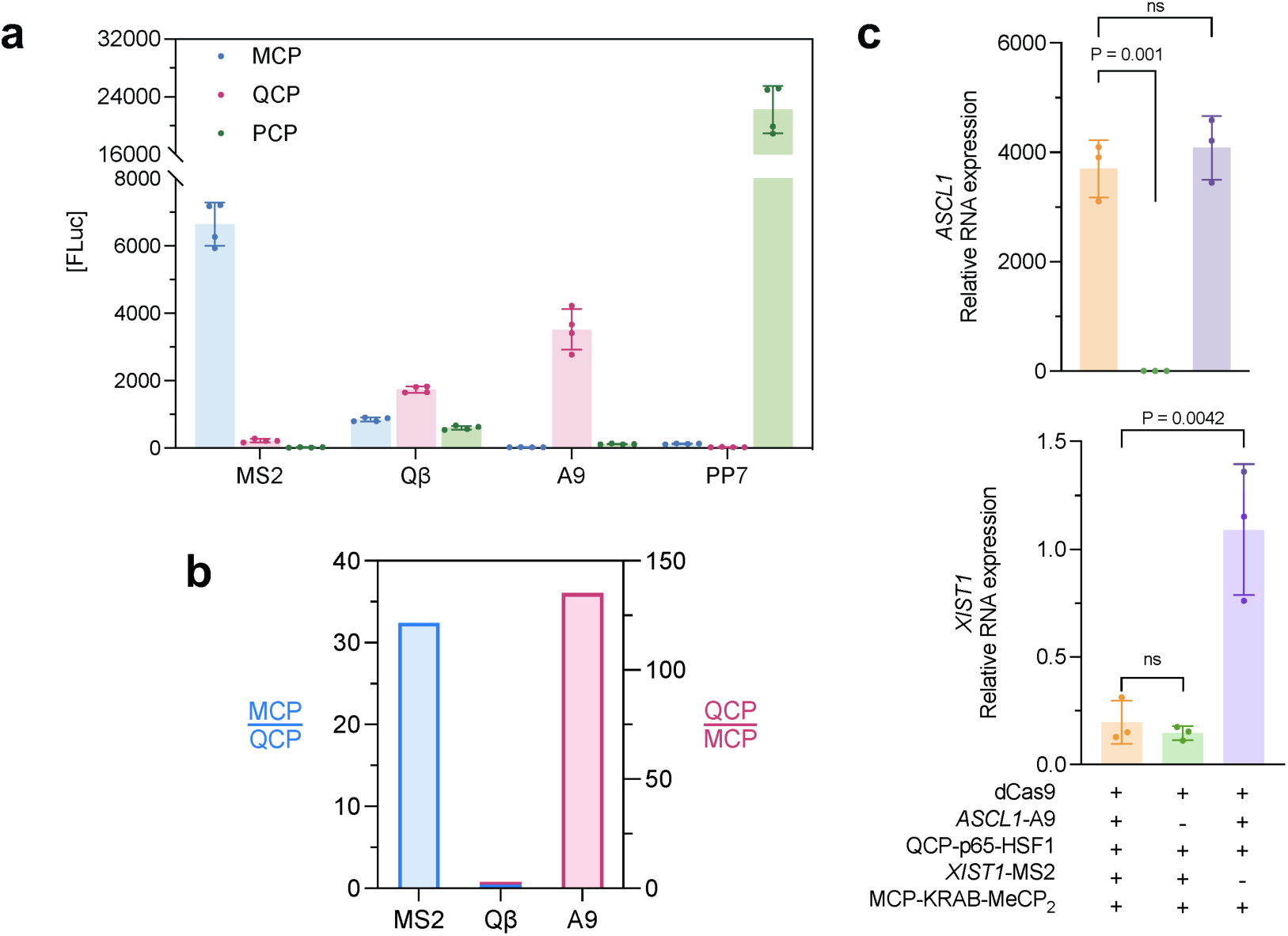
Orthogonal RNA-protein recruitments in HEK293T cells. **a**, No significant crossbinding is observed for MS2, PP7 and A9 with non-cognate RBPs, while evident crossbinding is observed for Qβ. (*n* = 4 technical replicates). **b**, A9 is highly specific for cognate target QCP over MCP, compared to Qβ and MS2. **c**, sgRNA-aptamer chimeras mediate simultaneously activation and repression at endogenous genes in HEK293T cells, measured by RT-qPCR. Fold of changes are calculated over cells missing both A9 and MS2 sgRNA chimeras. (*n* = 3 independent replicates). Values in **a** and **c** are mean ± s.d. ns, not significant. *P* values were determined by one-tailed paired Student’s *t*-test.

### Multifunctional transcriptional modulation in human cells

Next, we sought to explore the application of A9 in aptamer-mediated, multifunctional CRISPR transcriptional control in human cells. We designed multiplex sgRNA-A9 chimera targeting the *ASCL1* gene for activation with QCP-p65-HSF1, while another sgRNA-MS2 chimera simultaneously targeting the *XIST1* gene for repression with orthogonal MCP-KRAB-MeCP_2_ in HEK293T cells. We observed simultaneous significant activation of *ASCL1* (3,701-fold) and 88% repression of *XIST1* measured by RT-qPCR. These folds of change are similar to those observed when single genes are targeted by the respective activation or repression construct (**Fig. 4c**). Therefore, thanks to its orthogonality to existing aptamer-RBP pairs, the A9-QCP pair could be utilized in multiplexed CRISPR for simultaneous multifunctional modulation of endogenous genes.

## Discussion

Over the last ten years, a wide range of CRISPR technologies have emerged for gene manipulation, epigenetic functionalization, and transcriptional regulation. Among them, fusing effector proteins directly to the Cas protein allows the resulting CRISPR machinery to direct these effector proteins to multiple sites of the same gene or multiple genes at once^35,37,40^. Although they can be used to target multiple genetic loci simultaneously, these methods are often limited to applying one regulatory function (*e.g.*, activation or repression) at a time. On the other hand, recruiting effector proteins via aptamer-RBP recognition enabled multiplexed and multifunctional gene manipulations^44,48^. However, there are only a limited set of aptamer-RBP pairs that can function orthogonally and intracellularly, *e.g.*, MS2-MCP and PP7-PCP. The scarcity of orthogonal intracellular aptamer-RBP pairs imposes severe constraints on the CRISPR-mediated multifunctional manipulations of the genome and the epigenome. This work has expanded the scope of aptamer-RBP toolkit for CRISPR transcription regulators by establishing an intracellular directed evolution platform for sgRNA-fused aptamers. Applying this selection strategy, we successfully identified A9, a highly specific aptamer for QCP with an *in vitro* dissociation constant (*K*_d_) of 10 nM.

A major challenge with implementing aptamers identified by SELEX for *in vivo* applications is that the aptamers enriched from selections in simple buffers *in vitro* often fail to function properly *in vivo*, as a result of the non-specific interactions in the highly crowded and complex intracellular environment. For example, we found the previously reported QCP aptamer Qβ exhibited low intracellular activity and crossbinding with other proteins (**Fig. 4a**). To overcome this challenge, herein we directly evolve aptamers intracellularly using CRISPR-hybrid. Recognition of aptamer library by RBP of interest in a CRISPR-hybrid construct recruited the transcriptional activator and enhanced the expression of a GFP reporter, allowing cells carrying the functional aptamer hits to be enriched using FACS. We performed a mock selection with 1:1000 mixtures of cells carrying functional and non-functional aptamers, and observed >960-fold enrichment after two rounds of sorting. When applied to intracellular selection of aptamers from a diverse (>10^6^) library, the CRISPR-hybrid construct successfully enriched new aptamer sequences targeting bacteriophage coat proteins.

Orthogonality of the aptamer-RBP pairs is key to enable multiplexed and multifunctional manipulations of the genome and the epigenome. To improve the specificity of the evolved aptamer, we performed two intracellular selections of RNA libraries that originated from the MS2 aptamer and randomized the upper stem/loop region and the lower stem region, respectively. After the second selection, we observed top ten hits displayed significant lower crossbinding to MCP; and notably, we successfully obtained an aptamer, A9, that displays a stronger binding activity and preference to QCP than MCP. *In vitro* SPR affinity characterization confirmed the specificity of A9 to QCP. Importantly, although A9 exhibited modest affinity to MCP *in vitro*, it showed no crossbinding to MCP and PCP in a CRISPR-hybrid construct in mammalian cells, suggesting that the A9-QCP pair is orthogonal other known aptamer-RBP pairs in genome editing applications *in vivo*. To demonstrate the utility of orthogonal aptamer-RBP pairs in multiplexed CRISPR, we designed sgRNAs carrying A9 and MS2 to simultaneously activate and repress endogenous genes through the recruitment of QCP-p65-HSF1 activator and MCP-KRAB-MeCP_2_, respectively. Simultaneous transcriptional activation and repression at respective targeted genes was successfully achieved.

Beyond bacteriophage coat proteins, the intracellular selection system developed in this work can be used to discover aptamers capable of recruiting endogenous effector proteins, such as transcription factors, epigenetic editors and readers, kinases and phosphatases, DNA/RNA repair enzymes, translocation regulators, etc. Doing so would eliminate the need for the delivery of exogenous RBP-effector fusion proteins and allow CRISPR genomic and epigenomic editing system to recapitulate the endogenous processes and pathways more accurately. Furthermore, compared to genes encoding the exogenous fusion, DNA elements encoding aptamers are much smaller and easier to deliver by size-sensitive vectors such as adeno-associated virus (AAV).

In summary, we demonstrated that the intracellular aptamer evolution platform based on the CRISPR-hybrid construct is a promising strategy for discovering highly specific aptamers functional in a wide range of CRISPR-related technologies and other intracellular applications. The intracellular evolved aptamers are particularly suited for multiplexed and multifunctional editing of the genome and the epigenome. Going forward, the application of this approach to identify aptamers capable of recognizing and recruiting endogenous effector proteins will greatly expand our ability to interrogate complex networks of genes and their regulating proteins.

### Reagent availability

CRISPR-hybrid plasmids for library selection will be distributed by Addgene.

## Acknowledgements

The authors thank Mitchel Tepe and Zeiyi Huang for feedback on Python scripts for HTS analysis, Dr. Mi Zhou for assistance with mammalian cell culture, Weiqi Qiu for assistance with *in vitro* transcription, and Professor Abhishek Chatterjee for the helpful discussion. This work was supported by NIH Director’s New Innovator Award 1DP2HG011027-01 (J.N.) and National Science Foundation Graduate Research Fellowship Program (Q.S.-T.). Illustrations were created with BioRender.com.

## Authors contributions

Q.S.-T. and J.N. designed the project and prepared the manuscript. J.N. supervised the research. Q.S.-T. and J.F. conducted the experiments. K.F. and P.W. assisted with flow cytometry measurements and cell culture. K.F., P.W., and Y.H. assisted on molecular cloning and protein purification. H.R. edited the Python scripts. P.A. conducted FACS.

## Methods

### Molecular cloning

DNA oligos longer than 60 bases were purchased from Integrated DNA Technologies (IDT), while shorter oligos were purchased from Azenta Life Sciences. For CRISPR-hybrid selection, the MS2 aptamer in pCD061 (Addgene #113315) was removed to make pQS14.3 for aptamer library construction. Selection plasmids were modified from pCD185 (Addgene #113317). For mammalian cell assays, dCas9 was cloned from plasmid #156501 (Addgene), p65-HSF1 was cloned from plasmid #61423 (Addgene), and KRAB-MeCP_2_ was cloned from plasmid #167902 (Addgene). sgRNA-chimeras were modified from plasmid #61424 (Addgene). PCR was performed using Q5 High-Fidelity DNA polymerase from New England Biolabs (NEB). Genes were either synthesized as bacterial codon-optimized gBlocks fragments (IDT) or amplified by PCR from listed sources. All PCR products were purified using MinElute PCR Purification Kit (Qiagen) to 30 μL final volume and quantified using a NanoDrop™ One/One^C^ Microvolume UV-Vis Spectrophotometer (Thermo Scientific). Gibson Assembly was performed using NEBuilder HiFi DNA Assembly Master Mix (NEB) according to the manufacturer’s instructions. The hybridized constructs were transformed into electrocompetent *E. coli* Turbo cells (NEB) or TOP10 cells (ThermoFisher Scientific) according to the manufacturer’s instructions. Transformants were recovered in 1 mL Super Optimal broth with Catabolite repression (S.O.C.) medium for 1 h at 37°C, 250 r.p.m. Transformants were selected on Luria Bertani (LB) agar plates supplemented with appropriate antibiotic(s). Individual clones were inoculated in LB for overnight growth and plasmids were purified using GeneJET Plasmid Miniprep Kit (ThermoFisher Scientific) and verified by Sanger sequencing (Azenta Life Sciences).

### Luciferase assay in bacterial cell

Reporter plasmid was co-transformed with accessory and selection plasmids of interest into electrocompetent S1030 cells (Addgene #105063) and recovered using Davis rich media^60^ (DRM). Transformants were plated onto 1.8% agar-2x YT plates supplemented with antibiotics. After overnight growth at 37°C, single colonies were incubated in 2 mL DRM supplemented with antibiotics for 16 h at 37°C, 250 r.p.m. Cultures were diluted 1,000-fold in a 96-well deep well plate supplemented with antibiotics and inducers isopropyl-β-D-thiogalactoside (IPTG), or anhydrotetracycline (ATc) to induce protein expression when necessary. After growth for 5 h at 37°C, 200 µL of culture was transferred to a 96-well black wall, clear bottom plate (Costar), and the *A_600nm_* and luminescence for each well was measured on a plate reader. The *A_600nm_* of a well containing only media was subtracted from each sample to obtain a correct *A_600nm_* value. The raw luminescence value of each well was then divided by that well’s corrected *A_600nm_* value to obtain the luminescence value normalized to cell density. Each experiment was completed in at least biological triplicate, and the error bars shown reflect the standard deviations of the measurements. gRNA sequences are listed in Supplementary Table 1.

### Flow cytometry

*E. coli* MG1655 K12 cells (Addgene #37854) transformed with reporter, accessory and or selection plasmids were inoculated in 3 mL LB medium supplemented with appropriate antibiotics and grown for 17 h at 37°C, 250 r.p.m. Cells were pelleted by centrifugation for 5 min at 14,000 r.p.m., followed by one wash with phosphate-buffered saline, pH 7.4 (PBS) (Fisher Scientific).

Cells were then diluted 50-fold in PBS and analyzed on Becton Dickinson Accuri C6 Plus (BD Biosciences). To enrich for single cells, a side scatter threshold trigger (SSC-H) was applied. To gate for single bacterial cells, we first selected events that appeared on the center of the FSC-A vs. SSC-A plot, then selected events along the diagonal of the FSC-H vs FSC-A plot. Events that appeared on the edges of the fluorescence histogram were excluded.

### Cell culture

HEK293T cells (ATCC) were maintained at 37°C and 5% CO_2_ in DMEM-high glucose, GlutaMAX Supplement, pyruvate (Gibco) supplemented with 10% (v/v) heat-inactivated fetal bovine serum (Corning). All references to DMEM below refer to the complete medium described here.

### Transient gene expression

HEK293T cells were plated seeded approximately 500,000 cells per well in 6-well plates and transfected the next day. HEK293T cells were co-transfected with plasmids encoding dCas9-P2A-Puro (500 ng), a mix containing equal masses of sgRNA-targeting endogenous genes, QCP-p65-HSF1 (500ng), and MCP-KRAB-MeCP_2_ (500ng). Sequences of the sgRNA are provided in Supplementary Table 2. Transfection complexes were prepared using Lipofectamine 3000 (Life Technologies), following the manufacturer’s instructions. Cells were treated with 3.5 μg/ml puromycin at 24 h post-transfection. Media with puromycin was refreshed after 24 h, and cells were collected 72 h post-transfection.

### Quantitative RT-PCR

Cells were washed once with PBS, and total RNA was isolated with Trizol (Life Technologies), following the manufacture’s protocol. cDNA was synthesized using SuperScript III reverse transcriptase (Life Technologies), according to manufacturer’s protocol, priming from anchored oligo-dT_21_, random hexamers (Life Technologies). qRT-PCR was performed using Universal SYBR Green Master Mix (Abclonal) and gene specific primers (Supplementary Table). cDNA template-less was used as a negative control. Bulk gene expression measurements were normalized to a GAPDH internal control, and fold-changes were calculated against no sgRNA control group. qPCR primers are listed in Supplementary Table 3.

### Luciferase assays in mammalian cell

The sgRNA target sequence is listed in Supplementary Table 2. HEK293T cells were plated at approximately 2.0 x 10^4^ cells per well in a 96-well plate (ThermoFisher Scientific) and cultured for 24 h at 37°C in 5% CO_2_. The cells were then transfected with Lipofectamine 3000 (Invitrogen) in Opti-MEM (Gibco) according to the manufacturer’s protocol. Plasmids encoding for dCas9, RBP-p65-HSF1, sgRNA and luciferase reporter were transfected at a 1:1:1:1 ratio. The total amount of DNA was 0.1 μg per well. Bioluminescence measurements were obtained 48 h post-transfection using Dual-Glo Luciferase Kit (Promega) with plate reader.

## Directed evolution of RNA aptamers

### Library construction

The single-stranded DNA (ssDNA) libraries were synthesized by IDT, using a customized recipe (A: 25%, C: 25%, G: 25%, T: 25%) for random regions. Oligonucleotides are listed in Supplementary Table 4. ssDNA libraries were purified by 8% TBEU-Urea gel electrophoresis (Invitrogen) with RNA Clean & Concentrator kit (Zymo Research), and eluted with nuclease-free water. Plasmid pQS14.3, which only contains sgRNA scaffold without aptamer insertion, was used as template to generate aptamer libraries. Piece A was amplified with primers QS94 and QSL2 or QSL5 for library N11 and N10, respectively. Piece B was amplified with primers QS91 and QS92. Primerless overlap extension PCR (OEPCR) was performed in 16 x 50 µL reactions (10% DMSO, 20 cycle number) containing 100ng piece A and equimolar piece B. Piece C was amplified with primers QS95 and QS96. All PCR products were treated with *DpnI* (NEB) at 37°C for 2 h to digest any residual template plasmid, followed by purification on 1.5% TAE-agarose gels using QIAquick gel extraction kit (Qiagen). Aptamer libraries were constructed with piece C and OEPCR product (1:4 molar ratio) by Gibson Assembly and ethanol-precipitated with yeast-tRNA (Invitrogen) to transform into electrocompetent Top10 cells. Freshly electroporated cells were recovered in 10 mL S.O.C. medium for 1 h at 37°C, 250 r.p.m., followed by 17 h growth in 150 mL LB supplemented with carbenicillin (100 µg mL^-1^). More than 10^8^ transformants were obtained to ensure library coverage. Plasmid library was isolated with the ZymoPURE II Plasmid Midiprep Kit (Zymo Research) for following transformation into the appropriate MG1655 K12 (+ accessory and reporter plasmids) electrocompetent cells.

### Electrocompetent strain preparation

Electrocompetent *E. coli* MG1655 K12 cells were transformed with reporter plasmid and accessory plasmid of interest, depending on the target RNA-binding protein. Single clones were inoculated in 10 mL LB media supplemented with chloramphenicol (25 µg mL^-1^) and kanamycin (50 µg mL^-1^) for 16 h at 37°C, 250 r.p.m. Next day, culture was diluted 100-fold in 1 L Super Optimal Broth (SOB) media and grown under identical condition until it reached mid-log-phase (OD_600_ = 0.5-0.7). Cells were pelleted in three JA10 tubes (Nalgene) centrifuged at 6,000 r.p.m. for 15 min at 4°C. The media was immediately decanted and the interior of the tubes was wiped with a few Kimwipes (Kimberly-Clark) to remove residual media and salts. Each cell pellet was quickly resuspended in 15 mL of pre-chilled, sterile filtered 15% glycerol in MilliQ purified water using a serological pipette and combined into one tube with ∼300 mL of 15% glycerol. The cells were centrifuged and washed an additional two times. After the last wash, the interior of the tube was wiped with Kimwipes to remove residual glycerol solution. The pellet with resuspended in 1.5 mL 15% glycerol and split into 50 µL aliquots, which were flash-frozen using liquid N_2_ bath and quickly transferred to -80°C for storage. More than 10^8^ transformants is usually obtained to ensure aptamer library coverage. In addition, empty electrocompetent MG1655 K12 cells prepared by this method typically yielded 10^9^-10^10^ cfu per µg of plasmid DNA and enable simultaneous transformation of all three plasmids.

### Antibiotic selection

Selection plasmids with and without MS2 aptamer were separately transformed into MG1655 K12 (+ accessory MCP and reporter plasmids) electrocompetent cells, and selected on LB-agar plates supplemented with appropriate antibiotics. A single colony from these two cell types was inoculated in 3 mL LB for 17 h at 37°C, 250 r.p.m. After measuring culture’s optical density (OD_600_), cells were mixed in ratios of 1:1000 MS2 aptamer: no aptamer. A total of 10^7^ cells were challenged by plating them onto a 150 mm petri dish (Fisher Scientific) containing LB-agar, plasmid maintenance antibiotics (100 µg mL^-1^ carbenicillin and 25 µg mL^-1^ chloramphenicol), and a concentration of kanamycin pre-determined to be above the MIC of the MG1655 strain lacking aptamer component of the CRISPR-hybrid system. Plates were incubated at 37°C for 17-24 h and surviving colonies were harvested with 10 mL LB medium, centrifuged and plasmids were purified via miniprep. The enriched selection plasmids were extracted using 1% TAE-agarose gel and re-transformed into MG1655 K12 (+ accessory MCP and reporter plasmids) electrocompetent cells for next round.

### Fluorescence-activated cell sorting (FACS)

For mock selection, cells with and without MS2 aptamer were mixed as describe above. For library selections, plasmid library was transformed into the appropriate MG1655 K12 (+ accessory and reporter plasmids) electrocompetent cells and grown in 15 mL LB with maintenance antibiotics for 17 h at 37°C, 250 r.p.m. Cells were harvested by centrifugation for 5 min at 4°C, 14,000 r.p.m. Cells were washed once and diluted 50-fold in cold PBS (Fisher Scientific) to analyze and sort on Becton Dickinson FACSAria cell sorter (BD Biosciences). The sorting gate was set to include library members with fluorescence higher than control, which are cells lacking the aptamer component of CRISPR-hybrid. The top 0.007% (total of 1,500 cells) were collected in 1 mL S.O.C. medium. Collected cells were harvested at 14,000 r.p.m. for 5 min, and resuspended in 12 µL of PCR-Rescue Buffer (1 mM Triton X-100, 20 mM Tris-HCl, pH 8.0, 2 mM EDTA) for cell lysis. The reaction was incubated at 95°C for 3 min and cooled to 4°C. This reaction mixture was used as DNA template for 3 x 50 µL PCRs with primers QS91 and QS96 to amplify the region flanking the aptamer library. PCR products were subsequently purified on 1% TAE-agarose gels using QIAquick Gel Extraction Kit (Qiagen) and eluted with 25 µL nuclease-free water. The enriched aptamers were subcloned back into plasmid backbone with the Gibson assembly protocol described above. Top10 transformed cells were grown in 15 mL LB under the same growth conditions as above. This enriched library was transformed again into the appropriate MG1655 K12 (+ accessory and reporter plasmids) electrocompetent cells and re-challenged with the selection condition.

### Restriction enzyme digestion analysis

Selection plasmids or PCR products flanking the region of the aptamer library isolated from each round of mock selection were digested under the following conditions: 500ng DNA, 1 µL of rCutSmart buffer (NEB), 1 µL of appropriate restriction enzyme (NEB), up to 10 µL of nuclease-free water. The reactions were incubated in a thermal cycler at 37°C for 30 min, and halted by subsequent heat denaturation at 65°C for 20 min. DNA was analyzed on a 1% TAE-agarose gel stained with ethidium bromide (Invitrogen). The relative recovery of MS2 aptamers was quantified in ImageJ (imageJ.nih.gov)

### High-throughput sequencing and data processing

QS_seq18 and Qs_seq19 primers (see Supplementary Table 5 for sequences) containing adapters for Amplicon Sequencing were used for PCR amplification of enriched aptamer library plasmids. PCR products were purified by 2% TAE-agarose gel using the QIAquick gel extraction kit (Qiagen), and submitted to Azenta Life Sciences for Amplicon Sequencing. Processing of sequencing data was performed using Python script, available on GitHub. A Q-score filter was applied to analyze the quality scores of each read in the FASTQ files, and any reads below the specific threshold (Q-scores lower than 10) was discarded. A mismatch filter then used to compare each read with the expected sequence in the fixed regions of the sgRNA scaffold, skipping the aptamer pool region containing randomized sequences. The sequences in the randomized region were then extracted and collected in a comma-separated file, along with each read’s count. The most abundant sequences (listed in Supplementary Table 6) were individually validated by reporter assays. Aptamer structures were predicted using Mfold webserver.

## Surface plasmon resonance (SPR)

### RNA aptamer preparation

Aptamer containing 3’ 24-mer poly (A) sequence was purchased from IDT. RNA was purified with 8% TBE-Urea PAGE gel extraction (Invitrogen), recovered with crush and soak method, precipitated with ethanol, washed with 70% ethanol and dissolved in nuclease-free water. Purified RNA was verified once more by alkaline hydrolysis on a TBE-Urea PAGE gel (Invitrogen) stained with SYBR Gold dye (Invitrogen).

### Sensor chip surface generation

Experiments were performed on a OpenSPR (Nicoya) instrument at 23°C with HBS-EP+ (Cytiva) as the running buffer. Streptavidin (0.5 μM) was immobilized to ∼800 RU in both reference (FC1) and sample (FC2) flow cells on a biotin sensor chip using the Biotin-streptavidin Sensor Kit (Nicoya).

### Aptamer binding assay

Aptamers (∼1.5 μM, 10 μL) were diluted in running buffer for thermal treatment of heating 95°C for 3min and slow cooling to 4°C at 0.1°C s^-1^ in the MiniAmp thermal cycler (Thermo Scientific). A dilution series of protein analytes were prepared in running buffer and filtered through a 0.2 μm membrane (MilliporeSigma), using a minimum of six concentrations ranging from 0.1 x *K_D_* to 100 x *K_D_* for a more accurate estimate of the kinetic parameters. Each assay cycle includes a capture, association, dissociation, and regeneration step. The aptamer was further diluted to 0.1 μM in running buffer and captured onto FC2 for 1200 s at 5 μL/min. The target protein was injected over both FCs for 300 s at 30 μL/min, and running buffer was then injected over both FCs for 300 s to monitor target dissociation. Surface was regenerated with 25 mM of NaOH for 30 s at 30 μL/min over both FCs, which removed the captured RNA and protein. The TraceDrawer software (Nicoya) used to process the datasets and analyze interaction kinetics. The FC1 data was first subtracted from FC2 to correct for injection noise, baseline drift, nonspecific surface binding and bulk refractive index changes. This corrected data was fit to a 1:1 binding model.

### Bacterial expression and purification of recombinant proteins

BL21 (DE3) competent *E. coli* cells (NEB) transformed with plasmid pET22b-MCP-MBP-his_6_ or pET22b-Qβ-MBP-his_6_ was inoculated in 10 mL LB medium supplemented with carbenicillin (100 µg mL^-1^) for overnight culture at 37°C, 250 r.p.m. On Day 2, the culture was dilute 100-fold in 1 L LB medium supplemented with carbenicillin, and grown to OD_600_ of ∼0.5. Protein expression was induced with 1 mM final IPTG for 16 h at 16°C, 250 r.p.m. All purification steps were performed at 4°C with Stock Buffer (50 mM NaH_2_PO_4_ pH 8.0, 300 mM NaCl, 6 mM BME). Cells were harvested by centrifugation for 15 min at 6000 r.p.m, resuspended in 30 mL Lysis Buffer (Stock Buffer + 10 mM imidazole), and lysed by sonication (5 cycles of 20 s pulse-on, 1 min pulse-off at amplitude 80%). Lysate was homogenized by centrifugation at 14,000 rcf for 30 min. The clarified lysate was incubated with 1.5 mL Ni-NTA resin (Takara) for 1 h with gentle rotation and subsequently applied to an Econo-Pac^TM^ chromatography column (Bio-Rad Laboratories). The protein-bound resin was washed with 25 mL Wash Buffer (Stock Buffer + 50 mM imidazole), and His-tagged protein was eluted with 16 mL of Elution Buffer 1 (Stock Buffer + 100 mM imidazole) and 8 mL of Elution Buffer 2 (Stock Buffer + 200 mM imidazole). Fractions of 2 mL were collected and analyzed on 4-12% Bis-Tris PAGE gel (Invitrogen) with NuPAGE^TM^ MES SDS running buffer (Invitrogen) and Coomassie staining. Fractions without impurifies were pooled, buffer exchanged into Dialysis Buffer (Stock Buffer + 5% glycerol), and concentrated with a Amicon Ultra-15 Centrifugal Filter (MilliporeSigma), molecular weight cutoff of 10kDa, at 14,000 rcf for 15 min. Protein concentration was measured by Pierce™ Coomassie (Bradford) Protein Assay Kit (Thermo Scientific). Aliquots were flash-frozen at -80°C for storage or used directly for in vitro binding assay.

## References

1. Boch, J. et al. Breaking the code of DNA binding specificity of TAL-type III effectors. Science 326, 1509–1512 (2009).

2. Christian, M. et al. Targeting DNA Double-Strand Breaks with TAL Effector Nucleases. Genetics 186, 757–761 (2010).

3. Kim, Y. G., Cha, J. & Chandrasegaran, S. Hybrid restriction enzymes: zinc finger fusions to Fok I cleavage domain. Proc. Natl. Acad. Sci. 93, 1156–1160 (1996).

4. Bibikova, M., Golic, M., Golic, K. G. & Carroll, D. Targeted chromosomal cleavage and mutagenesis in Drosophila using zinc-finger nucleases. Genetics 161, 1169–1175 (2002).

5. Rouet, P., Smih, F. & Jasin, M. Expression of a site-specific endonuclease stimulates homologous recombination in mammalian cells. PNAS 91, 6064–6068 (1994).

6. Wang, H., La Russa, M. & Qi, L. S. CRISPR/Cas9 in Genome Editing and beyond. Annu. Rev. Biochem. 85, 227–264 (2016).

7. Cheng, C., Zhou, M., Su, Q., Steigmeyer, A. & Niu, J. Genome editor-directed in vivo library diversification. Cell Chem. Biol. 28, 1–10 (2021).

8. Strecker, J. et al. RNA-guided DNA insertion with CRISPR-associated transposases. Science 365, 48–53 (2019).

9. Anzalone, A. V et al. Search-and-replace genome editing without double-strand breaks or donor DNA. Nature 576, 149 (2019).

10. Woo Cho, S., Kim, S., Min Kim, J. & Kim, J.-S. Targeted genome engineering in human cells with the Cas9 RNA-guided endonuclease. Nat Biotech 31, 230–232 (2013).

11. Cong, L. et al. Multiplex Genome Engineering Using CRISPR/Cas Systems. Science 339, 819–823 (2013).

12. Gasiunas, G., Barrangou, R., Horvath, P. & Siksnys, V. Cas9-crRNA ribonucleoprotein complex mediates specific DNA cleavage for adaptive immunity in bacteria. PNAS 109, E2579–2586 (2012).

13. Jinek, M. et al. A Programmable Dual-RNA–Guided DNA Endonuclease in Adaptive Bacterial Immunity. Science 337, 816–821 (2012).

14. Mali, P. RNA-Guided Human Genome Engineering via Cas9. Science 339, 823–826 (2013).

15. Qi, L. S. et al. Repurposing CRISPR as an RNA-Guided Platform for Sequence-Specific Control of Gene Expression. Cell 152, 1173–1183 (2013).

16. Jinek, M. et al. RNA-programmed genome editing in human cells. Elife 2013, 1–9 (2013).

17. Hess, G. T. et al. Directed evolution using dCas9-targeted somatic hypermutation in mamallian cells. Nat. Methods 13, 1036–1042 (2016).

18. Komor, A. C., Kim, B., Packer, M. S., Zuris, J. A. & Liu, D. R. Programmable editing of a target base in genomic DNA without double-stranded DNA cleavage. Nature 533, 420– 424 (2016).

19. Nishida, K. et al. Targeted nucleotide editing using hybrid prokaryotic and vertebrate adaptive immune systems. Science 353, aaf8729-1–8 (2016).

20. Gaudelli, N. M. et al. Programmable base editing of T to G C in genomic DNA without DNA cleavage. Nature 551, 464–471 (2017).

21. Ma, Y. et al. Targeted AID-mediated mutagenesis (TAM) enables efficient genomic diversification in mammalian cells. Nat. Methods 13, 1029–1035 (2016).

22. Choi, J. et al. Precise genomic deletions using paired prime editing. Nat. Biotechnol. 40, 218–226 (2022).

23. Liu, X.-M., Zhou, J., Mao, Y., Ji, Q. & Qian, S.-B. Programmable RNA N 6-methyladenosine editing by CRISPR-Cas9 conjugates. Nat. Chem. Biol. 15, 865–871 (2019).

24. Wilson, C., Chen, P. J., Miao, Z. & Liu, D. R. Programmable m 6 A modification of cellular RNAs with a Cas13-directed methyltransferase. Nat. Biotechnol. 38, 1431–1440 (2020).

25. Thakore, P. I., Black, J. B., Hilton, I. B. & Gersbach, C. A. Editing the epigenome: technologies for programmable transcription and epigenetic modulation. Nat. Methods 13, 127–137 (2016).

26. Morita, S. et al. Targeted DNA demethylation in vivo using dCas9–peptide repeat and scFv–TET1 catalytic domain fusions. Nat. Biotechnol. 34, 1060–1065 (2016).

27. Pulecio, J., Verma, N., Mejía-Ramírez, E., Huangfu, D. & Raya, A. CRISPR/Cas9-Based Engineering of the Epigenome. Cell Stem Cell 21, 431–447 (2017).

28. Amabile, A. et al. Inheritable Silencing of Endogenous Genes by Hit-and-Run Targeted Epigenetic Editing. Cell 167, 219–232 (2016).

29. McDonald, J. I. et al. Reprogrammable CRISPR/Cas9-based system for inducing site-specific DNA methylation. Methods Tech. 5, 866–874 (2016).

30. Vojta, A. et al. Repurposing the CRISPR-Cas9 system for targeted DNA methylation. Nucleic Acids Res. 44, 5615–5628 (2016).

31. Xu, X. et al. A CRISPR-based approach for targeted DNA demethylation. Cell Discov. 2, 1–12 (2016).

32. Hilton, I. B. et al. Epigenome editing by a CRISPR-Cas9-based acetyltransferase activates genes from promoters and enhancers. Nat. Biotechnol. 33, 510–517 (2015).

33. Cheng, A. W. et al. Casilio: a versatile CRISPR-Cas9-Pumilio hybrid for gene regulation and genomic labeling. Cell Res. 26, 254–257 (2016).

34. Gilbert, L. A. et al. Genome-Scale CRISPR-Mediated Control of Gene Repression and Activation. Cell 159, 647–661 (2014).

35. Gilbert, L. A. et al. CRISPR-Mediated Modular RNA-Guided Regulation of Transcription in Eukaryotes. Cell 154, 442–451 (2013).

36. Bikard, D. et al. Programmable repression and activation of bacterial gene expression using an engineered CRISPR-Cas system. Nucleic Acids Res. 41, 7429–7437 (2013).

37. Perez-Pinera, P. et al. RNA-guided gene activation by CRISPR-Cas9–based transcription factors. Nat. Methods 10, 973–976 (2013).

38. Konermann, S. et al. Genome-scale transcriptional activation by an engineered CRISPR-Cas9 complex. Nature 517, 583–588 (2014).

39. Cheng, A. W. et al. Multiplexed activation of endogenous genes by CRISPR-on, an RNA-guided transcriptional activator system. Cell Res. 23, 1163–1171 (2013).

40. Chavez, A. et al. Highly efficient Cas9-mediated transcriptional programming. Nat. Methods 12, 326–328 (2015).

41. Nelles, D. A. et al. Programmable RNA Tracking in Live Cells with CRISPR/Cas9. Cell 165, 488–496 (2016).

42. Konermann, S. et al. Optical control of mammalian endogenous transcription and epigenetic states. Nature 500, 472–476 (2013).

43. Shechner, D. M., Hacisuleyman, E., Younger, S. T. & Rinn, J. L. Multiplexable, locus-specific targeting of long RNAs with CRISPR-Display. Nat. Methods 12, 664–670 (2015).

44. Zalatan, J. G. et al. Engineering Complex Synthetic Transcriptional Programs with CRISPR RNA Scaffolds. Cell 160, 339–350 (2015).

45. Dong, C., Fontana, J., Patel, A., Carothers, J. M. & Zalatan, J. G. Synthetic CRISPR-Cas gene activators for transcriptional reprogramming in bacteria. Nat. Commun. 9, 1–11 (2018).

46. Briner, A. E. et al. Guide RNA Functional Modules Direct Cas9 Activity and Orthogonality. Mol. Cell 56, 333–339 (2014).

47. Li, C. et al. SWISS: multiplexed orthogonal genome editing in plants with a Cas9 nickase and engineered CRISPR RNA scaffolds. Genome Biol. 21, 1–15 (2020).

48. Truong, V. A. et al. CRISPRai for simultaneous gene activation and inhibition to promote stem cell chondrogenesis and calvarial bone regeneration. Nucleic Acids Res. 47, (2019).

49. Ellington, A. D. & Szostak, J. W. In vitro selection of RNA molecules that bind specific ligands. Nature 346, 818–822 (1990).

50. Tuerk, C. & Gold, L. Systematic Evolution of Ligands by Exponential Enrichment: RNA Ligands to Bacteriophage T4 DNA Polymerase. Science 249, 505–510 (1990).

51. Wurster, S. E., Bida, J. P., Her, Y. F. & Maher, L. J. Characterization of anti-NF-κB RNA aptamer-binding specificity in vitro and in the yeast three-hybrid system. Nucleic Acids Res. 37, 6214–6224 (2009).

52. Wurster, S. E. & Maher, L. J. Selections that optimize RNA display in the yeast three-hybrid system. RNA vol. 16 253–258 (2010).

53. Hook, B., Bernstein, D., Zhang, B. & Wickens, M. RNA-protein interactions in the yeast three-hybrid system: Affinity, sensitivity, and enhanced library screening. RNA 11, 227– 233 (2005).

54. Sengupta, D. J. et al. A three-hybrid system to detect RNA-protein interactions in vivo. PNAS 93, 8496–8501 (1996).

55. Zhang, J. et al. Repurposing CRISPR/Cas to Discover SARS-CoV-2 Detecting and Neutralizing Aptamers. Adv. Sci. 2300656, 1–15 (2023).

56. Rumnieks, J. & Tars, K. Crystal structure of the bacteriophage qβ coat protein in complex with the rna operator of the replicase gene. J. Mol. Biol. 426, 1039–1049 (2014).

57. Witherell, G. W. & Uhlenbeck, O. C. Specific RNA Binding by Qß Coat Protein. Biochemistry 28, 71–76 (1989).

58. Horn, W. T., Tars, K. & Grahn, E. Structural Basis of RNA Binding Discrimination between Bacteriophages Qb and MS2. Structure 14, 487–495 (2006).

59. Lim, F. & Peabody, D. S. RNA recognition site of PP7 coat protein. NAR 30, 4138–4144 (2002).

60. Carlson, J. C., Badran, A. H., Guggiana-Nilo, D. A. & Liu, D. R. Negative selection and stringency modulation in phage-assisted continuous evolution. Nat. Chem. Biol. 10, 216– 222 (2014).

