## Supplementary Tables and Figures for "CRISPR-Hybrid: A CRISPR-Mediated Intracellular Directed Evolution Platform for RNA Aptamers"

**Table of Contents**

**Supplementary Tables**

Supplementary Table 1. gRNA sequences used in *E. coli* cells.

Supplementary Table 2. gRNA sequences used in HEK293T cells.

Supplementary Table 3. qPCR primers used in this work.

Supplementary Table 4. Amplification primers used to generate the aptamer libraries.

Supplementary Table 5. Oligonucleotides used for NGS.

Supplementary Table 6. Aptamers used in this work.

**Supplementary Figures**

Supplemental Figure 1: CRISPR-hybrid optimizations.

Supplemental Figure 2: Antibiotic mock selection with RpoZ activator.

Supplemental Figure 3: Optimizations with $SoxS_{R93A}$ transcriptional activator.

Supplemental Figure 4: Antibiotic mock selection with $SoxS_{R93A}$ activator

Supplemental Figure 5: FACS optimizations.

Supplemental Figure 6: Initial intracellular selection for QCP.

Supplemental Figure 7: A1 mutants binding activity and specificity.

Supplemental Figure 8: Second intracellular selection for QCP.

Supplemental Figure 9: SPR kinetic measurements.

**Supplementary Table 1. gRNA sequences used in *E. coli* cells.**

| **Name** | **Sequence (5’ to 3’)** |
|---|---|
| G0 | GUGAACCGCAUCGAGCUGAA |
| G1 | AACACGCACGGUGUUACAUU |
| G2 | UAGGGACCCUACAACACGCA |
| G3 | CGUUUUGCUGAGGAGACUUA |
| G4 | CUGGGCCUUUCGUUUUGCUG |
| G5 | AAAGGCUCAGUCGAAAGACU |
| G6 | GUAACACCGUGCGUGUUGUA |
| G7' | AGUCUCCUCAGCAAAACGAA |
| G8' | AGCCUUUCGUUUUAUUUGAC |
| G9' | AUGUCCUCUUGACAGUAAGA |
| G10' | AUCCGGUCAAAUAAAACGAA |
| G11' | CUUACUGUCAAGAGGACAUC |
| G12' | UUACCCGUCUUACUGUCAAG |
| 1.1 | AACACGCACGGUGUUACAUU |
| 1.2 | CGGAUCUUCCACAACACGCA |
| 1.3 | UUAUCACCUUGGCUGCAGGC |
| 1.4 | AUGUAACACCGUGCGUGUUG |
| 1.5 | CGUGCGUGUUGUGGAAGAUC |
| 1.6 | GAAGAUCCGGCCUGCAGCCA |

**Supplementary Table 2. gRNA sequences used in HEK293T cells.**

| Name | Sequence (5' to 3') |
|---|---|
| Gluc | GUGAAUGAUGAUAAUACGAUG |
| *ASCL1-1* | GCUGGGUGUCCCAUUGAAA |
| *ASCL1-2* | GCAGCCGCUCGCUGCAGCAG |
| *ASCL1-3* | GUGGAGAGUUUGCAAGGAGC |
| *ASCL1-4* | GUUUAUUCAGCCGGGAGUC |
| *XIST1-1* | GGCAGCGCUUUAAGAACUGAA |
| *XIST1-2* | GGACUGAAGAUCUCUCUGCACUU |
| *XIST1-3* | GGCCAUAUUUCUUACUCUCUCG |

**Supplementary Table 3. qPCR primers used in this work.**

| Target | Forward primer | Reverse primer |
|---|---|---|
| *ASCL1* | GGAGCTTCTCGACTTCACCA | AACGCCACTGACAAGAAAGC |
| *XIST1* | AGGTCAGGCAGAGGAAGTCA | CTGCCTCCCGATACAACAAT |
| *GAPDH* | TTCGACAGTCAGCCGCATCTTCTT | GCCCAATACGACCAAATCCGTTGA |

**Supplementary Table 4. Amplification primers used to generate the aptamer libraries**

| Oligo | Sequence (5' to 3') | Description |
|---|---|---|
| QSL2 | GTTAAAATAAGGCTAGTCCGTTATCAACTTGAAAAAGTGCGCACATNNNNNNNNNNNATGTGCTTTTTTTGAAGCTTGGGCCCGAACAAA | Forward primer for piece A of N11 library |
| QSL5 | TAAAATAAGGCTAGTCCGTTATCAACTTGAAAAAGTGCGCNNNNTAGCTAAGCTANNNNGCTTTTTTTGAAGCTTGGGCCCGAACAAAA | Forward primer for piece A of N8 library |
| QS94 | ACTCGAGTAAGGATCCAGTTCACCGAC | Reverse primer for piece A |
| QS91 | GAAAAGTGCCACCTGACGTCTAAGAAACCATTATT | Forward primer for piece B |
| QS92 | TTTTCAAGTTGATAACGGACTAGCCTTATT | Reverse primer for piece B |
| QS95 | GTCGGTGAACTGGATCCTTACTCGAGT | Forward primer for piece C |
| QS96 | AATAATGGTTTCTTAGACGTCAGGTGGCACTTTTC | Reverse primer for piece C |

**Supplementary Table 5. Oligonucleotides used for NGS**

| Oligo | Sequence (5' to 3') |
|---|---|
| QS_seq18 | ACACTCTTTCCCTACACGACGCTCTTCCGATCTTTGACAGCTAGCTCAGTC |
| QS_seq19 | GACTGGAGTTCAGACGTGTGCTCTTCCGATCTGTTTTTGTTCGGGCCCAAG |

Sequencing adapters are highlighted in yellow

**Supplementary Table 6. Aptamers used in this work.**

| Aptamer | Sequence (5' to 3') |
|---|---|
| MS2 | ACAUGAGGAUCACCCAUGU |
| QB | AUGCAUGUCUAAGACAGCAU |
| A1 | ACAUUAGCUAAGCUAAUGU |
| A2 | ACAUGUACUAAGUACAUGU |
| A3 | ACAUAGGCUAAGCCUAUGU |
| A4 | GGAUAGCUAAGCUACC |
| A5 | UGCUAGCUAAGCUAACA |
| A6 | UGUUAGCUAAGCUAUCA |
| A7 | AGUUAGCUAAGCUAAUU |
| A8 | UUUAGCUAAGCUAGGG |
| A9 | CCUUAGCUAAGCUAAUG |
| A10 | UGAUAGCUAAGCUACCA |
| A11 | UUCUAGCUAAGCUAGGA |
| A12 | GAGUAGCUAAGCUACCC |
| A13 | UUAUAGCUAAGCUAUAG |

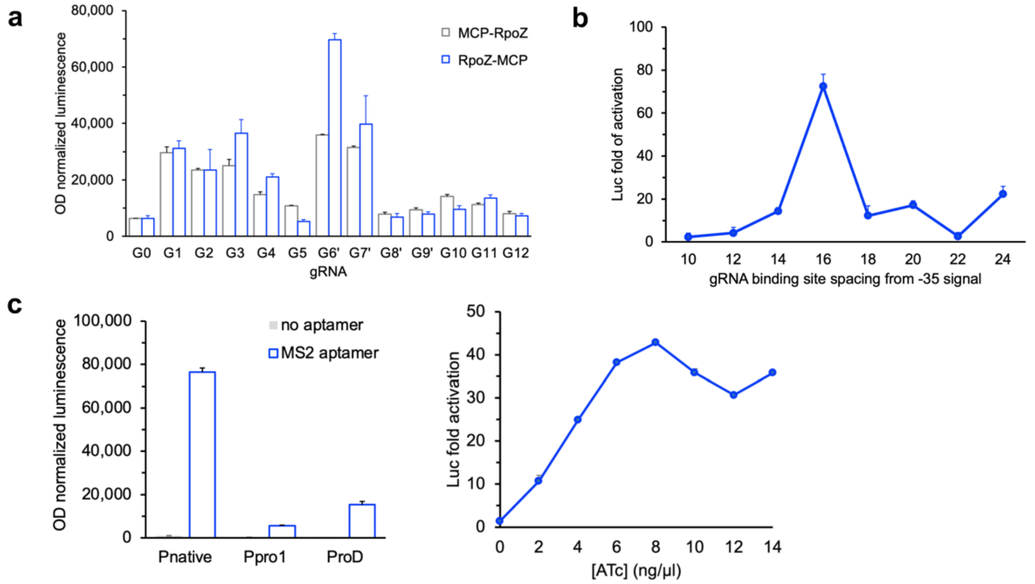


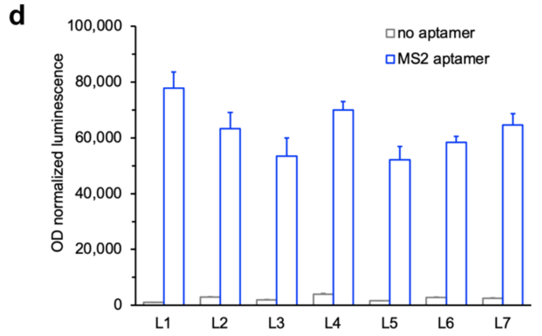


| Linker | Sequence | Length |
|---|---|---|
| L1 | AA | 2 |
| L2 | EQKLISEEDL | 10 |
| L3 | GGSGGSGGSGGSGGS | 15 |
| L4 | GHGTGSTGSGSS | 12 |
| L5 | MSRPDPA | 7 |
| L6 | GSAGSAAGSGEF | 12 |
| L7 | SGSETPGTSESATPES | 16 |

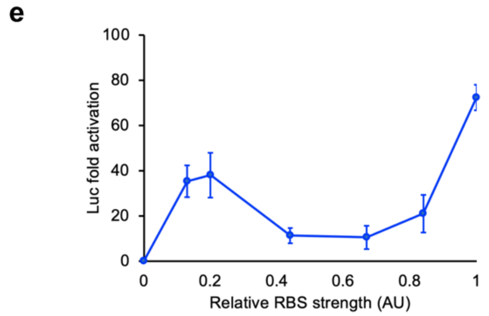


**Supplemental Figure 1: CRISPR-hybrid optimizations.** Luciferase expression in *E.coli* was used as reporter gene. **a**, highest levels of transcriptional activation were achieved with G6’ guide RNA and RpoZ-MCP. A scrambled guide RNA (G0) was used as negative control. **b**, G6’ binding at 16 base pairs upstream of the -35box in the J23107 promoter resulted in the most robust activation. **c**, Expression of dCas9 was driven by different constitutive promoters (left) and anhydrotetracycline inducible promoter (right). **d,** Tests of seven different linkers between MCP and RpoZ. **e**, Ribosome-binding site (RBS) modification enables robust modulation of the reporter activation levels. Data represent mean of three independent experiments ± s.d.

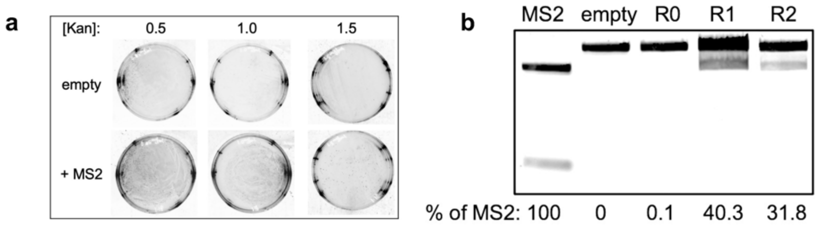


**Supplemental Figure 2: Antibiotic mock selection with RpoZ activator. a**, MS2 can recruit RpoZ-MCP to activate the expression of antibiotic resistance gene. e compared with *ScaI* and *AleI* digestion. Survival of cells harboring MS2 aptamer or no aptamer was measured on solid medium supplemented with increasing concentration of kanamycin. **b,** Gene compositions before and after mock selection were compared with *ScaI* and *AleI* digestion.

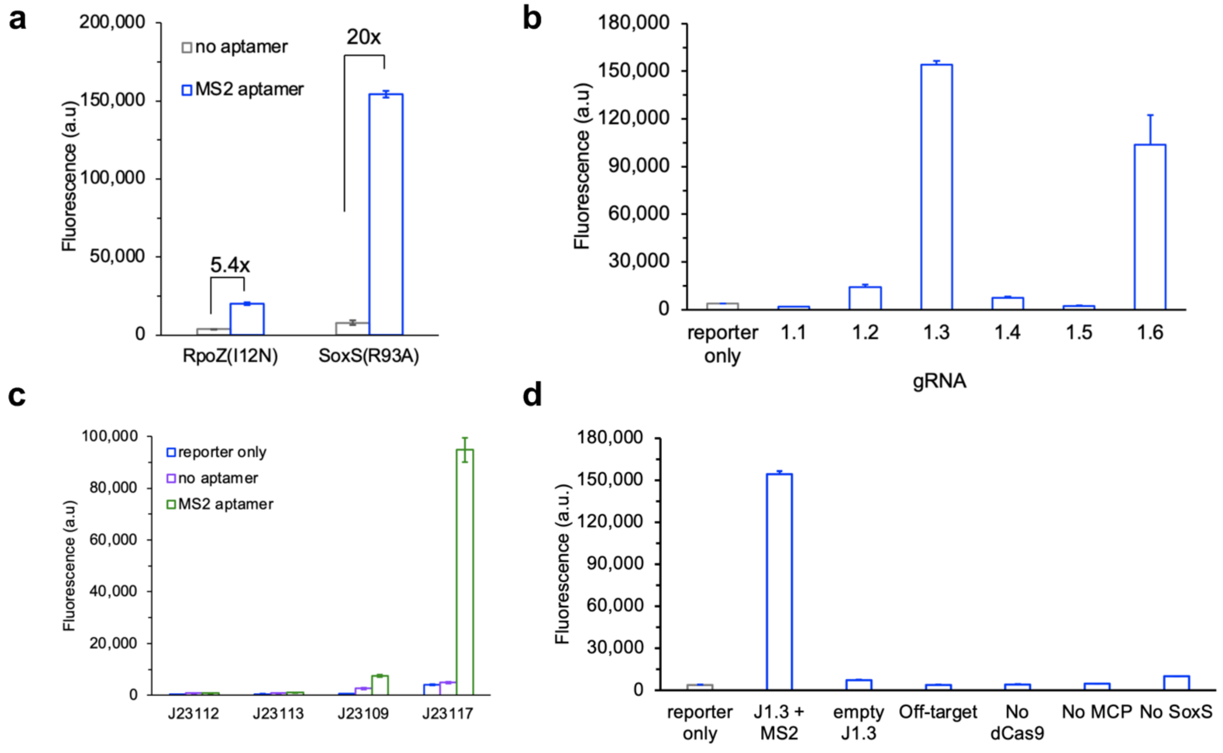


**Supplemental Figure 3: Optimizations with SoxS$_{R93A}$ transcriptional activator.** sfGFP was used as reporter gene. **a,** Transcriptional activator SoxS$_{R93A}$ showed 20-fold increase in reporter gene expression. **b,** Guide RNA 1.3 binding at 80 base pairs upstream of transcriptional start site yielded a 56-fold increase in GFP expression, compared to expression from reporter plasmid only. **c,** RNAP-minimal promoter affinity was evaluated with a series of bacterial J23 promoters, and J23117 resulted in the largest reporter fold activation by CRISPRa. **d,** Reporter activation was observed only when all components of CRISPRa are included. Data represent mean of three independent experiments ± s.d.

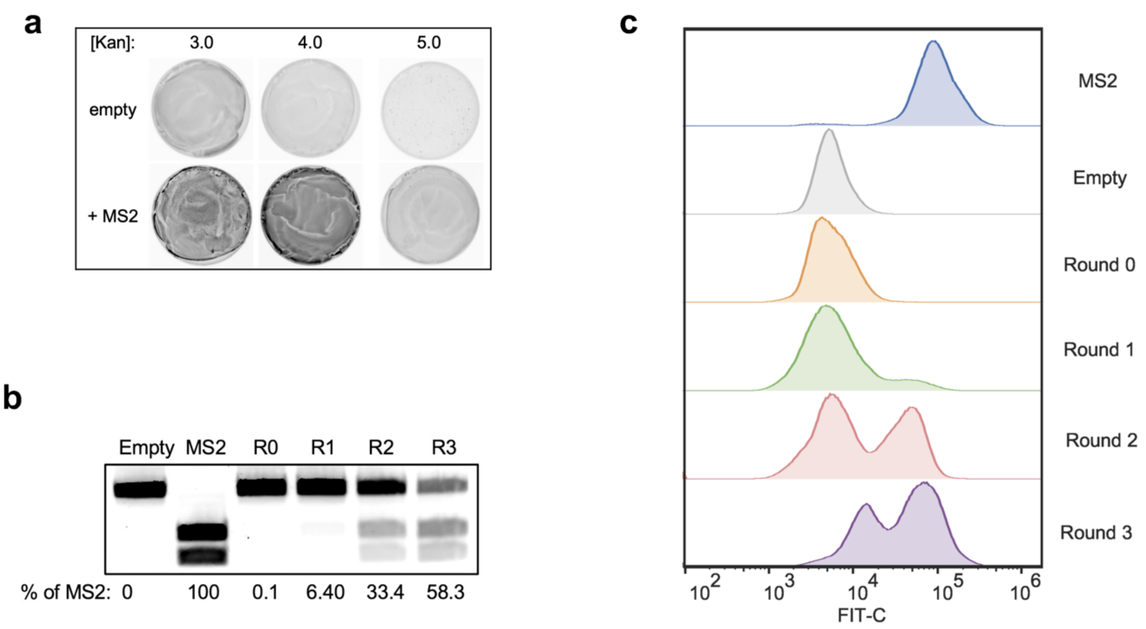


**Supplemental Figure 4: Antibiotic mock selection with SoxS$_{R93A}$ activator. a**, Dynamic range of [Kan-GFP] reporter. Survival of cells harboring MS2 aptamer or randomized aptamer was measured on solid medium supplemented with increasing concentration of kanamycin. Cells with MS2 recruiting MCP-SoxS$_{R93A}$ showed robust expression of Kan-GFP resistance gene for cell survival of up to 5.0mg/ml. **b**, Gene compositions before and after mock selection were compared with *ScaI* and *AleI* digestion. **c**, Fluorescence histogram overlays showed cell populations after antibiotic selection are distinguishable on flow cytometer.

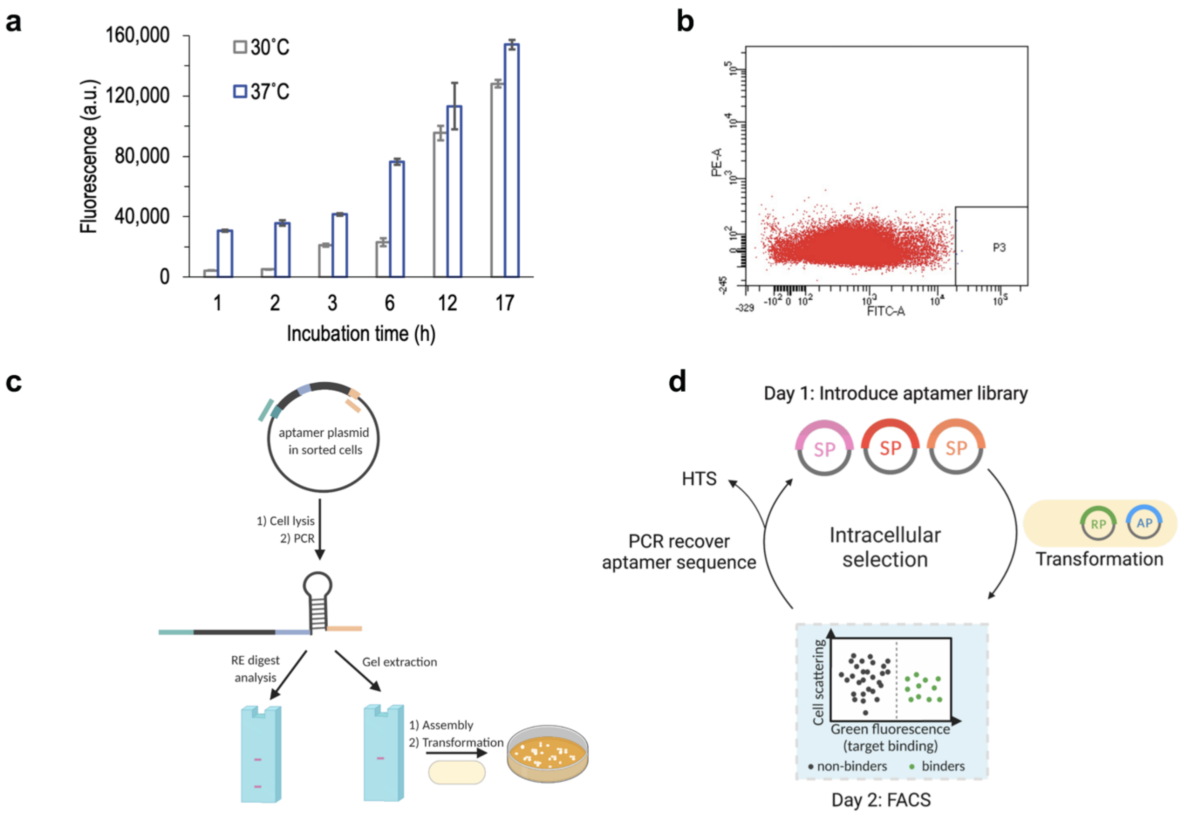


**Supplemental Figure 5: FACS optimizations. a**, Transcriptional activation of sfGFP reporter by CRISPR-hybrid with MS2 was measured at different *E.coli* growth temperature and time. Maximal sfGFP activation was observed from cells grown at 37°C for 17 h. Data represent mean of three independent experiments ± s.d. **b**, Representative plot showing gating strategy for FACS. Cells within the set gate (black rectangle) were collected. **c**, Schematic of the Rescue-PCR strategy for maximal recovery of sorted aptamer sequences. **d**, Overview of CRISPR-hybrid selection process coupled with FACS.

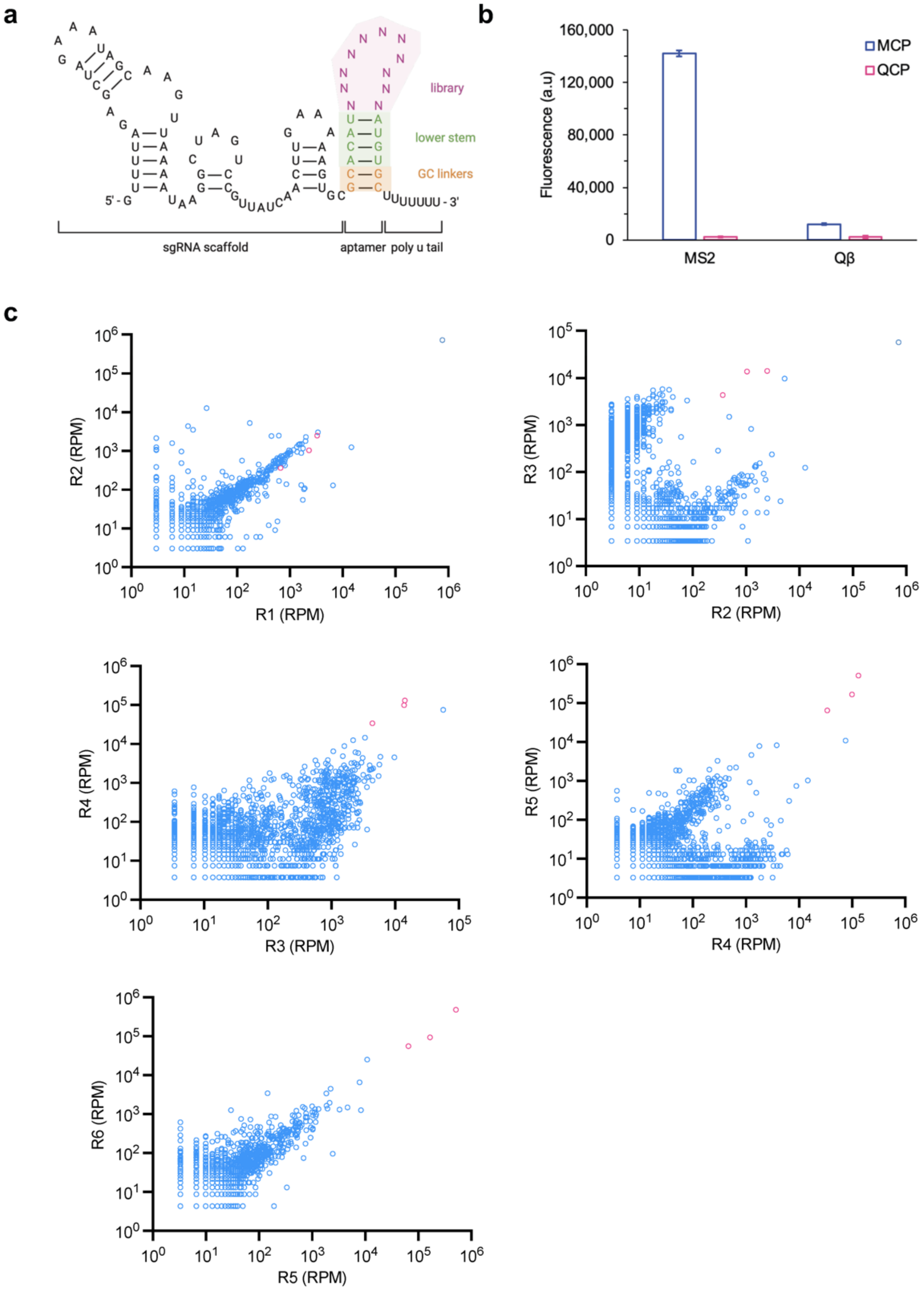


**Supplemental Figure 6: Initial intracellular selection for QCP. a,** Schematic of sgRNA scaffold with RNA aptamer library. **b,** Fluorescence measurements of GFP reporter activated by MS2 and QB binding to MCP and QCP. **c,** Scatter plots between two sequential rounds of library selection. Enrichment patterns observed are consistent with overlay histograms in Fig. 2c, and R5-R6 shows a high degree of correlation. The top three aptamers in R6 are colored in pink.

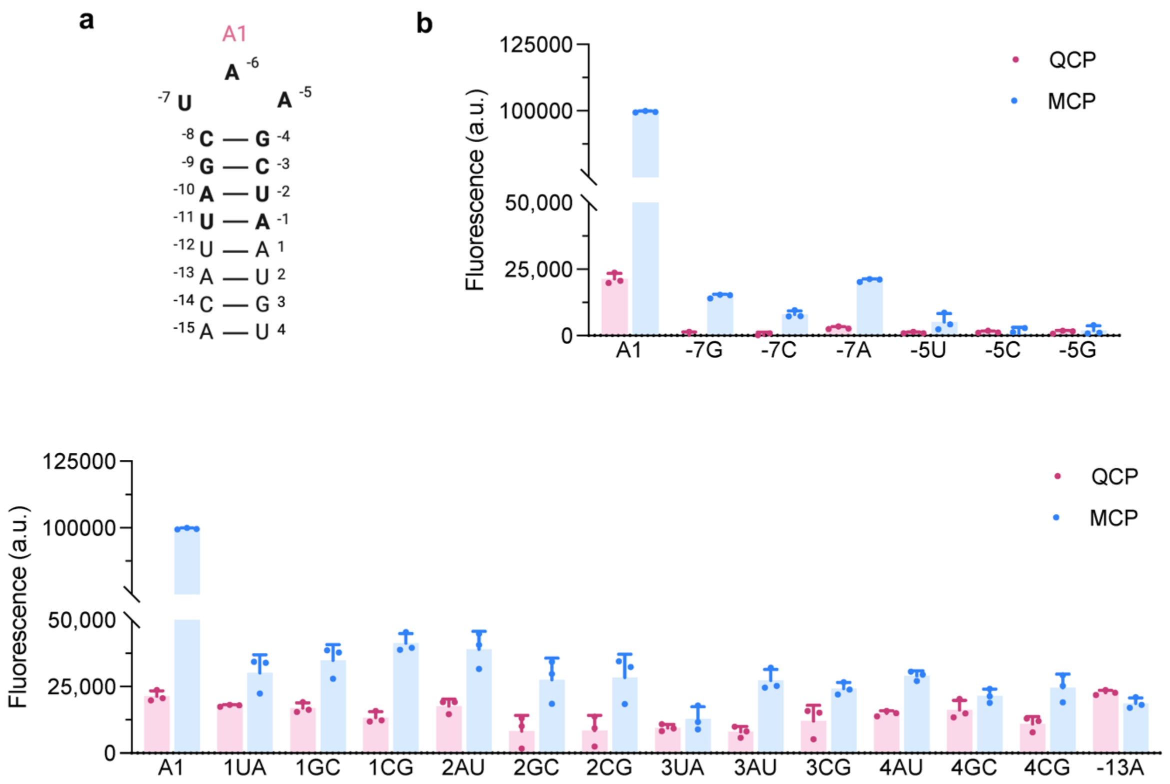


**Supplemental Figure 7: A1 mutants binding activity and specificity. a,** Structure of A1. **b,** Fluorescence measurements of A1 variants with single base mutations in the loop region. **c**, Fluorescence measurements of A1 variants with single base pair mutations in the lower stem region. Data represent mean of three independent experiments ± s.d.

a

b

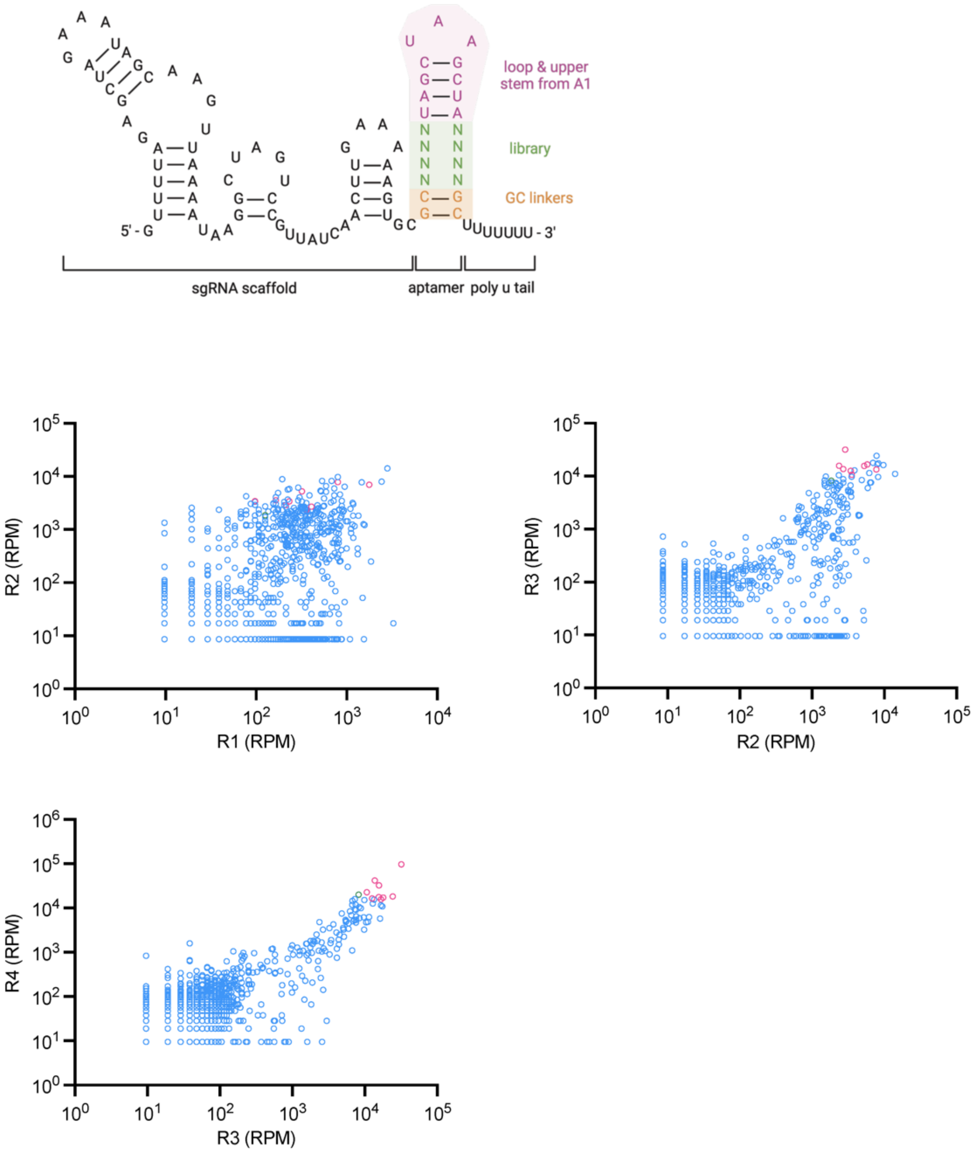


**Supplemental Figure 8: Second intracellular selection for QCP. a**, Schematic of sgRNA scaffold with RNA aptamer library. **b,** Scatter plots between two sequential rounds of library selection. Enrichment patterns observed are consistent with overlay histograms in Fig. 3a, and R3-R4 shows a high degree of correlation. The top ten aptamers in R4 are colored in pink with the most active aptamer, A9, colored in green.

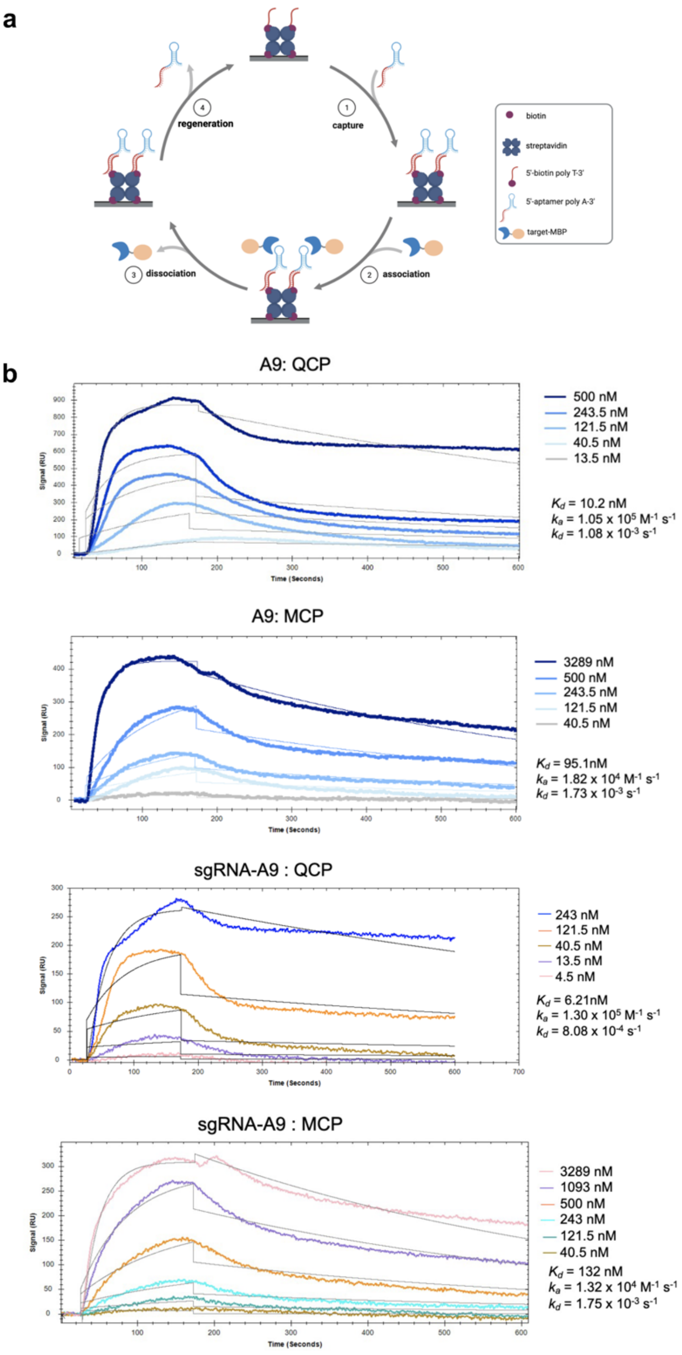


**Supplemental Figure 9: SPR kinetic measurements. a,** Illustration of SPR design. Biotinylated poly-T ssDNA is immobilized to streptavidin binding on biotin sensor. RNA aptamer extended with poly-A tail is captured onto the sensor by hybridizing with poly-T. Binding affinity of RNA aptamer with specific target protein is measured by recording the association and dissociation kinetics. Sensor is regenerated by removal of poly-A RNA aptamer for next cycle. **b,** Binding curves of A9 and sgRNA-A9 with QCP and MCP.
